# Intercalator effects on DNA loop extrusion by SMC proteins

**DOI:** 10.64898/2026.10.26.761753

**Authors:** Brian T. Analikwu, Jaco van der Torre, Cees Dekker

## Abstract

Intercalating dyes are widely used to visualize DNA in single-molecule experiments, but their effects on DNA-processing proteins remain incompletely characterized. Here, we study the effects of intercalators on DNA loop extrusion by SMC complexes. Intercalating dyes become fluorescent upon DNA binding, enabling direct visualization of extruded DNA loops. As dye intercalation alters the biophysical properties of DNA by lengthening and unwinding it, we examine how that modifies loop extrusion. We systematically characterize the effects of intercalators, and find that they stiffen DNA, partly neutralize its charge, and can substantially reduce the binding and ATPase activity of SMCs. We assess four dyes and conclude that SYBR Gold and SYTOX Orange are less invasive intercalators than SYTOX Green or YOYO-1. While loop extrusion is qualitatively not affected by intercalation, quantitative values for SMC binding and looping rates differ significantly from those in the absence of intercalators. We present a new dye-free loop-extrusion assay (utilizing local fluorescent spots along DNA), and find that loop extrusion proceeds at 1.75±0.26 kb/s for *S. cerevisiae* condensin without dyes, while adding intercalators leads to a two-fold lower rate. Our findings establish that intercalators quantitatively modify loop extrusion parameters, and we offer practical recommendations for minimizing their impact.

**Graphical abstract:** 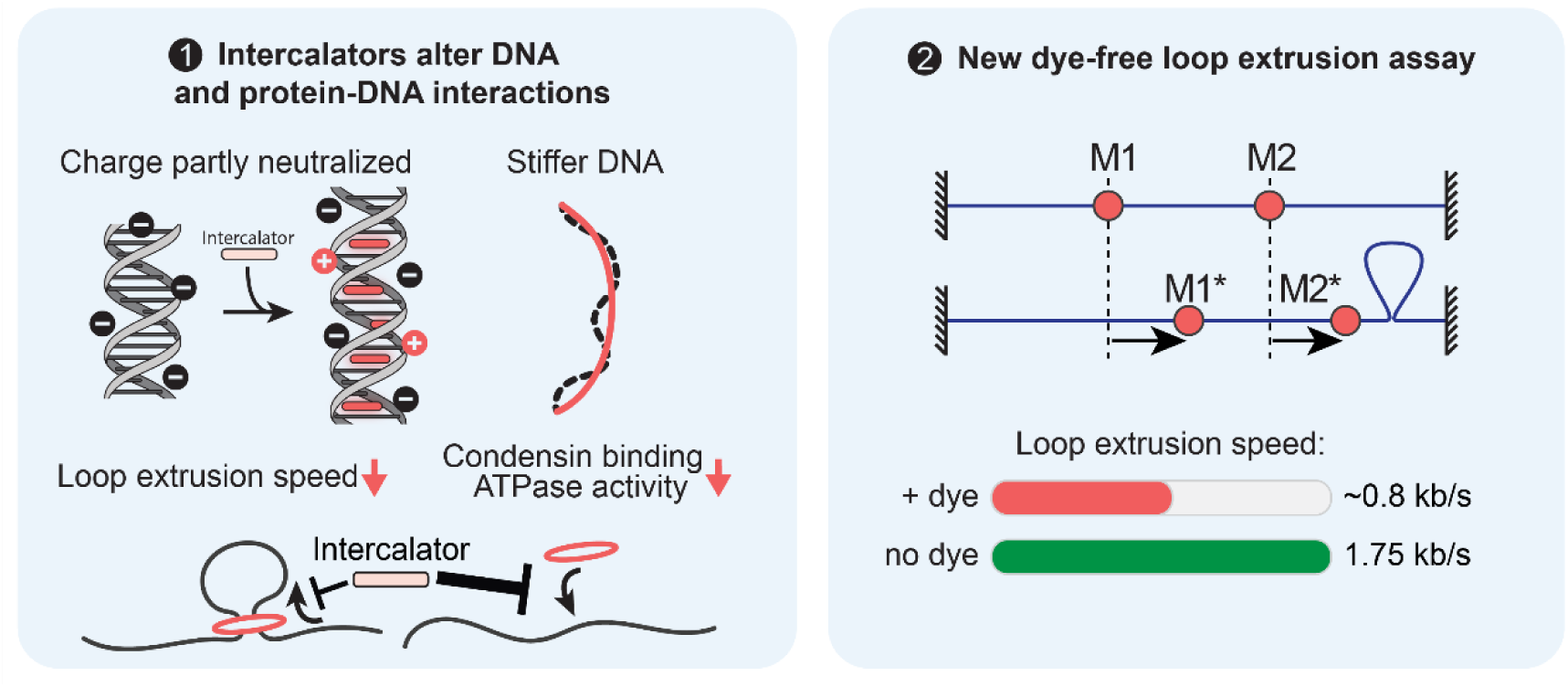

## Introduction

Intercalating dyes such as ethidium bromide, SYTOX Orange, and SYBR gold are widely used in biochemistry applications because of their unique ability to fluoresce upon binding to DNA. These dyes bind DNA in a specific binding mode as they are stacked between neighboring basepairs, first termed ‘intercalation’ in the early 1960s^1^. Ethidium bromide, the archetypical intercalating dye, was shown by the late 1960s to have significantly enhanced fluorescence upon intercalation^2,3^. Within a few years, this photophysical property was exploited to enable direct visualization of nucleic acids that were separated by gel electrophoresis^4^, establishing a nowadays routine technique in biochemical laboratories. Apart from their specific use in gel electrophoresis visualization, intercalator dyes are widely applied in e.g. the quantification of nucleic acids, monitoring of real-time and quantitative PCR, cell cycle analysis in flow cytometry, as well as in *in vitro* visualization of single molecules of nucleic acids. While microscopy technology improved significantly since then, the first single molecules of DNA were already visualized with DNA-binding dyes in the early 1980s^5,6^, where their characteristic dynamic fluctuations became apparent.

It is well established that intercalating dyes alter the DNA mechanics. Their binding extends the DNA length, it induces twist by locally unwinding the helix, it alters the DNA flexibility, and positive charges on the dyes partly neutralize the DNA’s negative phosphate backbone^1,7,8^. The length increase is rather well understood. A distinction can be made between mono-and bis-intercalators, where the former insert a single planar region between stacked nucleotide bases, whereas the latter insert two planar regions between two close-by sets of neighboring basepairs. Upon intercalating, a mono-intercalator therefore extends the DNA by roughly the distance between basepairs, whereas a bis-intercalator by roughly double that value (though the exact value depends on the specific dye and measurement used)^1,8,9^. A less clear picture exists for the change in stiffness of DNA, which is typically expressed as the persistence length. Some reports indicated a strong increase^10–13^ or decrease^14,15^ of the persistence length upon intercalation, while others reported no changes at all^8,16–19^, even for the same dyes. A potential cause for this discrepancy is the high stretching force on DNA that is often used to determine the persistence length using magnetic or optical tweezers. Since intercalators exhibit different binding parameters (affinity, size of the binding site) as a function of DNA tension^8,14,20^, it is likely that this resulted in different stiffness values. While it is clear that intercalator binding affects the length, twist, stiffness, and overall charge of the DNA molecules under study, these changes are often underestimated in experimental studies.

There are many fluorescence-based single-molecule assays in which DNA is necessarily visualized. These assays include DNA curtains^21^, optical tweezers^22^, and single-molecule assays where linearized DNA is visualized by TIRF/HILO as it is attached to a coverslip at either one or both ends, typically mildly stretched by flow^23^. The main advantage of the latter assay over DNA curtains and optical tweezers is, respectively, the uniform tension profile and low tension on the DNA (with forces as low as <0.1 pN). This assay has recently been applied to a wide class of studies on DNA and DNA-binding proteins, including those on loop extrusion by SMC complexes (Structural Maintenance of Chromosomes)^24–26^, DNA binding and condensation by ParB proteins^27^, positioning and translocation of RNA polymerase^28^, nuclease dependence on supercoiling^29^, and many more. In all of these applications, intercalating dyes were used simply as a tool to visualize DNA while studying protein-DNA interactions. Yet, because intercalators do modify DNA’s mechanics, it is important to understand how these dyes affect the protein-DNA interactions of interest. While a few notable efforts have been made to study the effect of intercalators on DNA-binding proteins^8,29–32^, it remains unclear which of the dye-induced changes to DNA underlie these effects.

Here, we investigate how intercalating dyes affect DNA processing by a specific class of DNA-binding proteins: SMC complexes, and in particular the condensin SMC complex from *S. cerevisiae*. These SMC complexes have recently emerged as the central chromosome-shaping proteins as they structure the 3D genome by generating large DNA loops, up to megabases in size, through a process called DNA loop extrusion. From the onset of mitosis, condensin acts to rapidly compact chromatin by extruding DNA loops in preparation for cell division^33^, while in interphase the cohesin SMC complex defines topologically associated domains in chromosomes. Strikingly, SMC complexes operate at the sub-piconewton force scale, making them much weaker molecular motors than other well-characterized motors such as kinesin^34^ or RNA polymerase^35,36^. The single-molecule assay in which DNA molecules are mildly stretched over a glass coverslip is therefore most suitable to study how this class of protein complexes interacts with DNA. Indeed, this assay was successfully applied to identify condensin, cohesin, and SMC5/6 as loop-extruding molecular motors ^24–26^, with many follow-up discoveries about their motor properties^32,37–41^. Moreover, this assay is exceptionally suited to study how these complexes interact with other DNA-binding proteins that they encounter as they linearly translocate along the DNA^37–39,41^. From these studies, we know that loop extrusion proceeds at a rate of 0.5-1.5 kb/s along bare DNA, can generate loops of many tens of kilobases before stalling due to tension, and typically does not get blocked by most roadblocks, with the rare exception for specific inhibitors like CTCF or very long dense stretches of proteins^24,37–39^. Notably, however, all these results were obtained in the presence of intercalating dyes on the DNA, which may indeed alter DNA binding and loop extrusion quantitatively.

Accordingly, we here set out to measure the impact of intercalator binding on DNA loop extrusion by the condensin SMC complex. To investigate the mechanism of how intercalators affect these properties, we first take a closer look at the biophysical properties of a panel of four intercalating dyes that are commonly used: YOYO-1, SYBR Gold, SYTOX Orange, and SYTOX Green. Despite their abundant usage in single-molecule assays, the latter two are relatively poorly characterized and have only very recently been structurally determined^42^. We first measure the relative amount of intercalator binding to obtain estimates for their binding affinity in relevant buffer conditions, which allows us to directly calculate the intercalator density (i.e., the number of intercalators per basepair). Next, we quantify the persistence length of DNA upon intercalation in the <0.1 pN tension regime to obtain a reliable measure for it in the relevant context for loop extrusion. We also quantify the electrostatic effects of intercalators binding to DNA. Furthermore, we characterize the intercalator effects on condensin-DNA interactions, such as on the bulk ATPase activity and on condensin binding to DNA. At the single-molecule level, we develop a new dye-free assay to directly obtain a measure for the loop extrusion rate and maximum loop sizes in the absence of any intercalator dyes. Together, the experiments link intercalator binding to changes in DNA biophysics, condensin function, and DNA loop extrusion, providing a framework for interpreting and minimizing the effects of DNA staining in single-molecule studies of chromosome organization.

## Materials and Methods

### Key resources

| Name | Source | Catalog number |
| --- | --- | --- |
| SYBR Gold | Invitrogen | S11494 |
| YOYO-1 | Invitrogen | Y3601 |
| SYTOX Orange | Invitrogen | S11368 |
| SYTOX Green | Invitrogen | S7020 |
| *S. cerevisiae* condensin | Ref.<sup>24</sup> | N/A |
| *E. coli* LacI<sup>SNAP-AF647</sup> | Ref.<sup>38</sup> | N/A |
| 50 kb DNA constructs | This work | N/A |
| Streptavidin | NEB | N7021S |
| Ultrapure BSA | Invitrogen/FisherScientific | AM2616 |
| PBS tablets | Sigma | P4417 |
| (±)-6-Hydroxy-2,5,7,8-tetramethylchromane-2-carboxylic acid (Trolox) | Sigma | 238813 |
| Malachite Green<br>Phosphate Assay Kit | Sigma | MAK307 |
| Disposable folded capillary cell | Sysmex / Malvern Panalytical | DTS1070 |

### Buffers

The concentration of SYBR Gold stock was determined on serial dilution and absorption measurement at 486 nm with an extinction coefficient of 57,000 M^-1^cm^-1^. Buffer solutions were prepared freshly before use. PBS (Phosphate buffered saline) was prepared from tablets. Standard LE buffer (loop extrusion buffer) consisted of 50 mM KGlu, 2.5 mM MgCl_2_, 20 mM Tris-HCl pH 7.5, 1 mM TCEP, 2 mM ATP, and 0.25 mg/ml BSA. The standard LE buffer was supplemented with other components depending on the assay as indicated in the relevant methods section. Intercalating dyes were stored in DMSO, but they were first sufficiently (>1000-fold) diluted in Millipore water, to dilute out DMSO before use.

### Characterisation of intercalator binding

The binding parameters of intercalators were determined using a titration and measuring the fluorescence as in ref.9. Corning 384-well plates (3575) were used in a Biotek Synergy H1 platereader set to 24°C. Filter settings (excitation/emission) were as follows: 495/537 for SYBR Gold, 480/521 for YOYO-1, 493/534 for SYTOX Green, and 545/571 for SYTOX Orange. 80 bp long oligonucleotides were used at a concentration of 1 nM, yielding an 80 nM basepair concentration. Intercalator concentration ranging from 0 to 5 μM (or 10 μM for YOYO-1) were used. The SYBR Gold stock concentration was determined from the absorbance as in ref.9.

Intercalator binding is then calculated using the same method as in ref.9 from fitting the McGhee-von Hippel equation to the observed fluorescence intensities *I*:

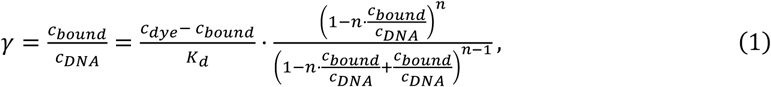

where *c_bound_* is the unknown concentration of bound dyes, which scales proportionally to the measured fluorescence intensity. Fitting was done using custom written code (see supporting code supplement).

### 50 kb DNA construct preparation

DNA constructs were made as described previously in ref.^43^. Briefly, we cloned plasmid pBeloBacIIggb-C0-2×4LacO-C10 (JT387) using our ModDNA toolkit, and made a 50 kb DNA construct with biotin handles from it. The plasmid DNA was first treated with RecBCD for 30 min at 37°C, followed by heat inactivation at 60°C for 10 min. The ModDNA-BAC plasmids were then cut with BsaI-v2 (NEB), heat inactivated at 65°C for 10 minutes and subsequently a ligation reaction was set up with the correct handles. Handles were duplex oligonucleotides carrying a 5’ phosphate and a 3’ biotin, with the complementary strand either unmodified or carrying an additional 5’ biotin (IDT). This gives one construct end a single biotin and the other end two biotins, which makes unconstrained biotinylated constructs. An approximate 10 to 100-fold excess of oligonucleotides was used and incubation was done overnight at 16°C with T4 DNA ligase (NEB). After ligation, a purification was performed using an S500HR gel filtration column (Cytiva). The procedure was conducted with an ÄKTA-start at room temperature. Alternatively for the construct with fluorescent handles, the handles were made by a PCR of 250nt, where during the PCR we had 1/5 Atto643-x-dUTP (Jenabioscience), and one primer a 5’ biotin. The PCR products were digested with BsaI-v2, purified using a PCR-cleanup kit from NEB and ligated instead of the duplex-oligonucleotides.

### Flow channel preparation

Flow channels for single-molecule loop extrusion assays and persistence length measurements were prepared as previously extensively described^23^ with some modifications. Briefly, borosilicate glass slides with laser-drilled holes and borosilicate coverslips were successively treated with acetone, 1M KOH, and acid piranha before extensive washing with anhydrous methanol. Aminosilanisation was then performed using 8% ATPMS ((3-aminopropyl)trimethoxysilane) in the presence of 4% glacial acetic acid in anhydrous methanol. The silanisation reaction was performed for 30-45min at room temperature followed by thorough rinsing with anhydrous methanol and drying at 70°C for at least 30 min. The slides and coverslips were then PEGylated using 1:40 Biotin-PEG5000-SVA : mPEG5000-SVA (Laysan Bio) in PEGylation buffer (either 100 mM borate buffer pH 8.5 or 100 mM NaHCO_3_/Na_2_CO_3_ pH 8.5). Slides and coverslips were incubated in a humidity chamber for at least 3h at 4°C followed by thorough rinsing with Millipore water and drying under N_2_-flow. This PEGylation was performed for successive rounds and final PEGylation was performed using MS(PEG)_4_-NHS in PEGylation buffer. Slides and coverslips were then assembled into flow chambers using laser-cut double-sided tape to mark the walls of the channels. Tubing on the outlet side was added and connected to a 1 mL syringe or syringe pump. Flow channels were finally incubated with 5 mg/ml Ultrapure BSA (Invitrogen), 10 μg/ml Streptavidin in 40 mM Tris-HCl pH 8.0, 20 mM NaCl buffer. The channel was incubated for >15 min before a brief rinsing step with LE buffer to remove unbound BSA and Streptavidin and to equilibrate the flow channel for the experiment.

### Single-molecule microscopy

The microscopy slides were positioned on a homebuilt HILO/TIRF microscope. Typically, intercalated DNA was visualized with a 0.4-0.8 mW 488 nm laser, though SYTOX Orange-bound DNA was visualized using a 0.5 mW 561 nm laser. LacI-dimers and fluorescent handles were visualized using a 30 mW 640 nm laser. Images were recorded with a PrimeBSI camera set to 100 ms per frame over a 1400×1400 FOV, resulting in an exposure time of roughly 83 ms.

### Persistence length measurement from lateral fluctuations

The persistence length of DNA molecules was measured from analysing the lateral fluctuations of individual DNA molecules tethered to a PEGylated glass surface. To this end, 50 kb DNA molecules were introduced into the flow channel at a flow rate of 2-5 μL/min using a syringe pump (New Era). Once a sufficient number of DNA molecules was bound in the flow channel, the buffer was exchanged to LE buffer without DNA and the flow was stopped abruptly by stopping the pump and closing a connecting valve. At least 1 minute of recording was made per condition, resulting in over 600 timepoints per individual DNA molecule. Recorded DNA molecules were then selected if they were sufficiently spatially separated from other DNA molecules to allow proper tracking. Only those molecules that remained intact throughout the recording (i.e. did not break due to photodamage) were used. The selected molecules were then cropped out of the total FOV and median-filtered to remove background signal. These background-filtered images were then used as input to a custom-written analysis pipeline (see supporting code supplement). In this pipeline, the tether endpoints were first identified globally by averaging all recorded frames, applying Otsu thresholding, and subsequent skeletonization. Each DNA molecule was then traced in each recorded frame. The traces were then smoothed using B-Spline interpolation over 200 points. The smoothed traces were then used as input to the bending-mode decomposition as introduced in the Results section and further detailed in Supporting Notes 1. After extracting the bending mode coefficients for each molecule at each timepoint along with the tension in each molecule according to the wormlike-chain model^44^, these coefficients and tensions were used to globally fit the persistence length per condition. DNA molecules for which the coefficients *a_k_* had an autocorrelation time > 100ms (i.e. the time per frame) were used. A worse autocorrelation indicated poor tracking. A correction for increased contour length due to intercalation was taken into account in these calculations, depending on the concentration and specific intercalator used. Error bars in Fig. 1E were generated from bootstrapping over DNA molecules.

**Figure 1.**
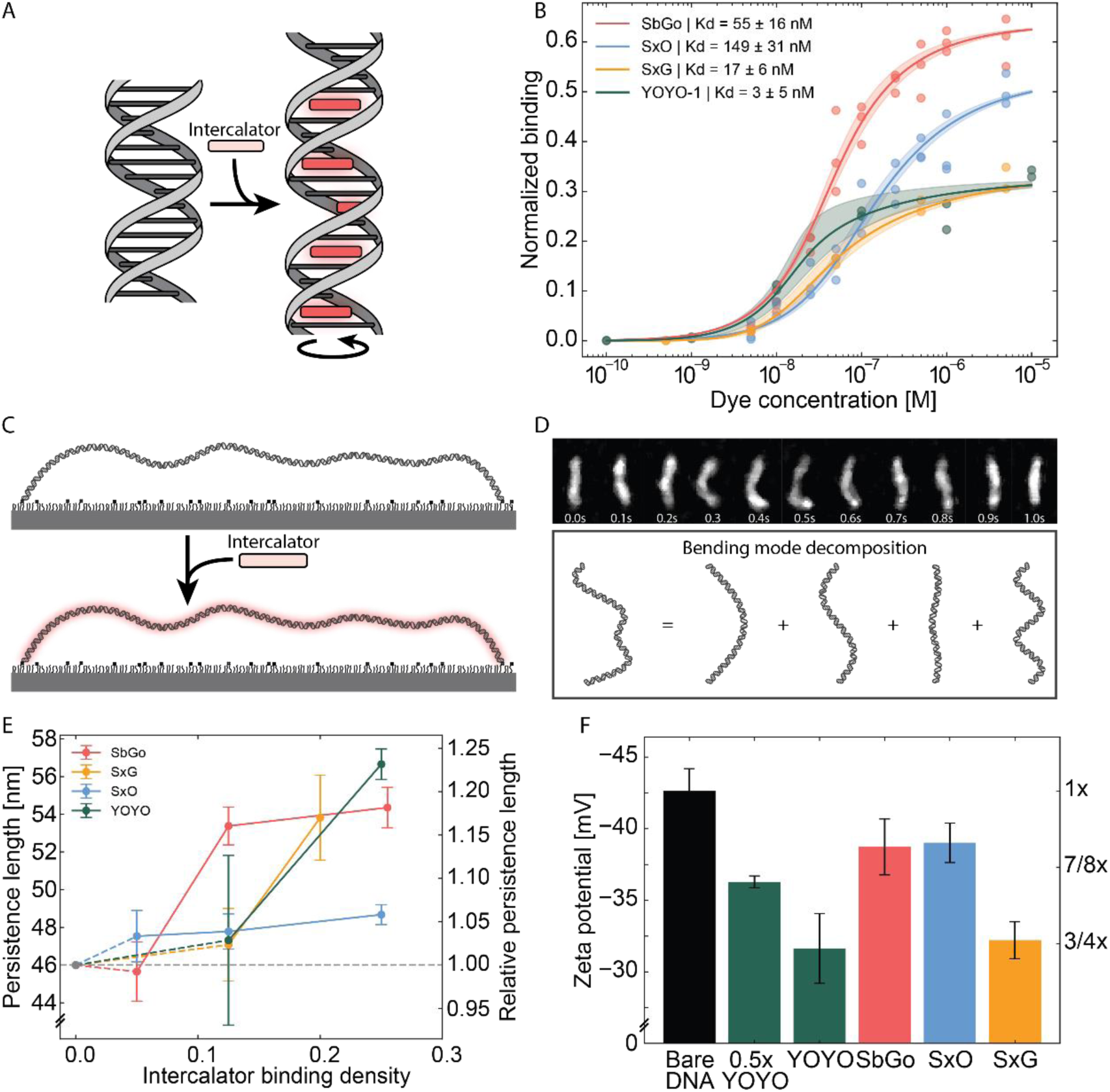
Characterisation of intercalator binding and its effects on DNA mechanics. **A)** Schematic representation of intercalation. The intercalator fluoresces upon binding between basepairs, thereby twisting and lengthening the DNA. **B)** Titration curves for SYBR Gold (SbGo), SYTOX Orange (SxO), SYTOX Green (SxG), and YOYO-1 dyes in loop extrusion buffer. Fitted affinities from the McGhee-von Hippel fits are given in the legend as fit ±standard deviation. Individual measurements are shown as dots, fitted curves are shown as lines, and standard deviation of the fits is indicated as the shaded area. **C)** Schematic of DNA bound on a glass slide. The glass is PEGylated with 40:1 PEG:Biotin-PEG for specifically binding biotinylated DNA through streptavidin. Intercalators are added to visualise DNA by fluorescence. **D)** 50 kb DNA observed over time (top), where the lateral fluctuations are related to the stiffness of the polymer, which can be identified by decomposing the tethered DNA molecule into half-harmonic bending modes. **E)** Persistence length measurement from lateral fluctuations in the presence of intercalators. Intercalator binding density is calculated from the McGhee-von Hippel fits in b. Bootstrap 95% CI are indicated as error bars. Apparent persistence length is given in nm (left y-axis) and as relative to the persistence length of DNA in the absence of intercalating dyes (46 nm, right y-axis). **F)** Zeta potential measurements on DNA in the presence of intercalator dyes. Bare DNA serves as a reference (“1x” on the right vertical axis). Intercalator was added at 1:4 dye:bp ratio except for the bar marked with ‘0.5x YOYO’, which had a 1:8 dye:bp ratio.

### Zeta potential measurements

To determine the charge of intercalated DNA, we used a Malvern Panalytical Zetasizer ZEN3600 and appropriate folded capillary zeta cell cuvettes (DTS1070). Typically, 800 μL samples were prepared. 200 ng/μl salmon sperm DNA was used, resulting in a 308 μM basepair concentration. Accordingly, 75 μM intercalator was added, resulting in a 4:1 bp:intercalator ratio. All experiments were conducted in a low-ionic buffer of 2 mM Tris-acetate pH 7.5. The peak zeta potential was extracted from each measurement and represented in Fig. 1F. The error bars report on the standard deviation of three replicates.

### Protein purification

*S. cerevisiae* condensin was purified as previously originally detailed and further clarified in refs.^24,38^. Fluorescently labelled *E. coli* LacI was purified as described in ref.^38^. Briefly summarized, full length LacI with a C-terminal SNAP-tag, followed by an HRV-3C protease cleavage site, and a His8-tag was expressed with a plasmid derived from pBAD, expressed in ER2566 cells (NEB). Cells were grown at 37°C in the presence of 100 μg/ml ampicillin and expression was induced with 0.2% (w/v) arabinose. Cells were then harvested after 3 hours of expression, followed by a PBS wash and resuspension. Lysis was performed using a CF1 cell disruptor at 20 kpsi at 4°C. After clarification, the lysate was applied to 2 mL Talon Superflow resin (Cytiva 28-9575-02) and incubated for 1h while rolling at 4°C. The resin was then washed with 50 mL PBS supplemented with 20 mM imidazole and 500 mM NaCl, followed by elution with 15 mL PBS supplemented with 200 mM imidazole. Proteins were then concentrated, labelled at room temperature with 100 μM SNAP-Surface Alexa Fluor 647 (NEB S9136S). Labelled LacI^SNAP-AF647^ was then purified further by size exclusion chromatography on an S200 Increase 10/300 column in 20 mM Tris-HCl pH 7, 100 mM NaCl, 50 μM TCEP.

### ATPase assay

The ATPase assay was performed using end-point measurements based on the Malachite Green Phosphate Assay Kit (MAK307, Sigma). *S. cerevisiae* condensin at 25 nM was incubated for 10 min at 30°C together with either 25 nM (Fig. 2E) or 1 μM 80 bp DNA (Fig. S5) and with the indicated amount of intercalator. Care was taken to dilute intercalator stocks such that the DMSO concentration never got above 0.5% as we observed that too high DMSO concentrations inhibited the ATPase activity by itself (Fig. S5). Incubations were halted by the addition of ice-cold EDTA solution to 10 mM. Malachite working buffer was added to the reactions and measurements of the produced phosphate were taken on a BioSPX Agilent Synergy H1 plate reader in absorption mode set to 620 nm with pathlength correction at 977/900 nm. For every measurement, a phosphate standard was included and samples were normalized against an intercalator-free condition with condensin, DNA, and ATP, while samples were blanked against intercalator-free LE buffer. This procedure yields a relative ATPase activity, with the intercalator-free LE condition set to 100%. Absolute quantification of the ATPase activity in this condition resulted in 4.4 ATP/s for *S. cerevisiae* condensin at 30°C.

**Figure 2.**
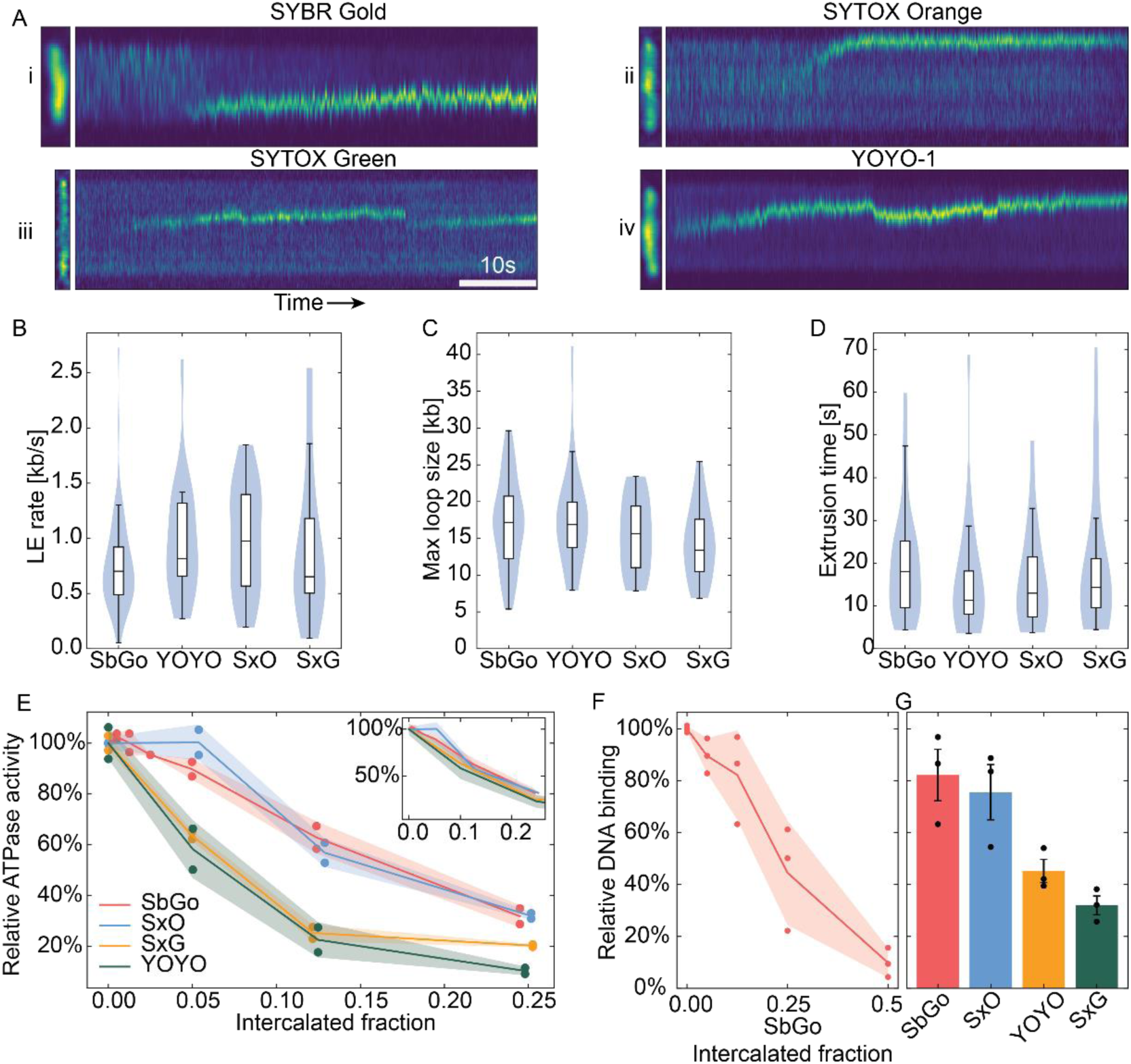
Intercalation inhibits DNA loop extrusion. **A)** 60s kymographs of loops formed in the presence of intercalating dyes at an intercalator density of 10%. **B-D)** Quantification of loop extrusion properties in the presence of different intercalating dyes. Loop extrusion rates, maximum loop sizes, and loop extrusion times, respectively. The large overlap in distributions and few significant differences indicate the similarity of these dyes with respect to their effects on loop extrusion. No significant differences were identified between any of the pairs of distributions (Kolmogorov-Smirnov p>0.5 for any pair). N= 58, 22, 15, and 19 respectively. **E)** ATPase activity in the presence of intercalating dyes at various binding densities. The shaded area represents the standard deviation. ATPase activity is normalized to the activity in the absence of intercalators. The strong decrease indicates inhibition of ATPase activity. Inset: The same data, but the x-axis represents SYBR Gold equivalent intercalated fraction, where the YOYO-1 and SYTOX Green axes are scaled by their ∼2-fold greater charge neutralization (from zeta-potential measurements). All lines collapse on a single curve. **F)** Condensin DNA binding in the presence of SYBR Gold at various binding densities, measured by EMSA. The shaded area represents the standard deviation. **G)** Same as F but for different dyes at a fixed intercalated fraction of 0.125. The error bars represent the standard error of the mean.

### Electrophoretic Mobility Shift Assay

To determine the effects of intercalation on the DNA binding ability of condensin, an EMSA was performed. 3% Agarose gels were prepared with 0.5X TBE. The gel and running buffer were pre-cooled on ice, while running was also performed on ice. 10 nM of condensin protein was added to 5 nM of 42 bp oligonucleotide dsDNA labelled with Cy5. Binding incubation was performed for 10min at room temperature in LE buffer supplemented with 1% PEG8000. After the incubation, glycerol was added to 5% to aid sample loading and the samples were immediately loaded on the gel. The gel was run at 90V for 20min and the gel was imaged on a Typhoon gel imager in the appropriate Cy5 channel. The fraction of condensin-bound DNA relative to the total DNA was then calculated from band intensity after background subtraction. For each replicate and intercalator used, the 0 nM intercalator condition was used as a reference for normalization.

### Mass photometry assay

Mass photometry was carried out on a Refeyn One mass photometer. Glass coverslips were cleaned by sonication in 1 M KOH, followed by rinsing with Millipore water. To allow DNA binding, the coverslips were incubated with 0.01% (w/v) poly-L-lysine (Sigma Merck) for 10 min before another rinse with Millipore water and blow-drying with N2 gas. 10 μL samples were prepared and incubated for 30 s to allow binding. 40 nM of 500 bp NoLimits (Thermo Scientific) and 10 nM *S. cerevisiae* condensin were incubated in LE buffer without BSA. Masses were calibrated using the NativeMark unstained protein standard (Invitrogen). At least three replicates were generated. Ensemble averages were generated over all replicates, followed by the generation of a kernel density estimate with a bandwidth of 20 kDa. From this kernel density estimate, Gaussian peaks were fitted around known positions (150, 510, 638, 840, 1276, 1550 kDa) with a distance up to 20 kDa away from these peak centres. Peak quantifications were then used to quantify condensin binding in Fig. S7. To obtain a confidence interval for the bound fraction, the delta method was applied. Because the bound fraction is a ratio of stoichiometry-weighted peak area, per-replicate ratios were not averaged directly, as this is biased when a replicate’s denominator is small (the mean-of-ratios problem). Each replicate was instead fit independently to obtain its DNA-bound and total condensin peak areas (bound_i, total_i); the reported estimate is the ratio of replicate means, R = bound/total (a combined ratio estimator). Its standard error was obtained via the delta method for a ratio of two correlated variables: 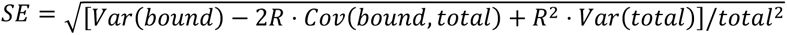. Single-shot K_d_ estimates were obtained assuming independent 1:1 binding based on Ref. ^45^ and reported in Fig. S7. Confidence intervals on the K_d_ estimates were obtained by evaluating at the lower and upper bounds of bound fractions.

### Loop extrusion assay

DNA was introduced in the flow cells as described above for the persistence length measurements. Instead of stopping the flow and imaging, condensin was introduced in the flow channel in the presence of LE buffer, supplemented with an oxygen scavenger (2.5% glucose, 18.75 nM glucose oxidase, 10 nM catalase, 1 mM Trolox activated to >10%). 20 mM Trolox ((±)-6-Hydroxy-2,5,7,8-tetramethylchromane-2-carboxylic acid) stock solution was prepared in 400 mM Tris-HCl pH 7.5, followed by activation with a 254 nm UV lamp (CAMAG, 022.9160). The condensin concentration was tuned in order to observe a sufficient number of loops for analysis, ranging from 20 pM in the absence of intercalator (Fig. 4) to 500 pM for the higher intercalator concentrations. DNA molecules in the presence of condensin were imaged for at least 10min in the absence of any flow, similar to the persistence length measurements described above. DNA molecules were selected based on the presence of a loop. Cropping of individual molecules and median-filtering for background suppression were also applied before analysis.

### Marker-modified loop extrusion assay

The marker-modified loop extrusion assay was performed similarly as described above, with the exception that DNA constructs at ∼5 pM were first incubated with 6 nM LacI^SNAP-AF647^. Incubation was performed for 10min at room temperature in LE buffer supplemented with 50 mM KGlu additionally, reaching 100 mM KGlu total. After the incubation, the DNA was introduced in the flow channel and the experiment was conducted identically as described above. If the experiment was performed with both the spotlabel markers and intercalating dye, the dye was added after the initial LacI incubation. DNA molecule selection was based on the presence of four spot labels (two at the ends and two at the LacO repeats) that were aligned in the direction of the flow. Measurements of spot locations were done in the absence of flow. When a loop was extruded, the LacI-specific spots in the middle moved jointly.

### Data analysis of loop extrusion

For the analysis of extruded loops based on the intercalated DNA signal, similar Python software was used as previously described^26,38,46^. This software allows the generation of kymographs and tracking of intensities and the sizes of loops based on the DNA signal. The same software with slight modifications was used to analyse extruded loops based on the movement of the spot labels. The extruded loop size was calculated as graphically illustrated in Fig. 3B and more examples are given in Fig. S8. First, the position of the LacI spotlabels was normalized between the two edges (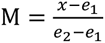, with *e_1_* and *e_2_* the positions of the edges, *x* the position of a marker, and M the normalized marker position). Second, the loop size was calculated based on the direction of the positional shift of the marker:

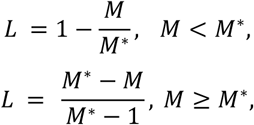

**Figure 3.**
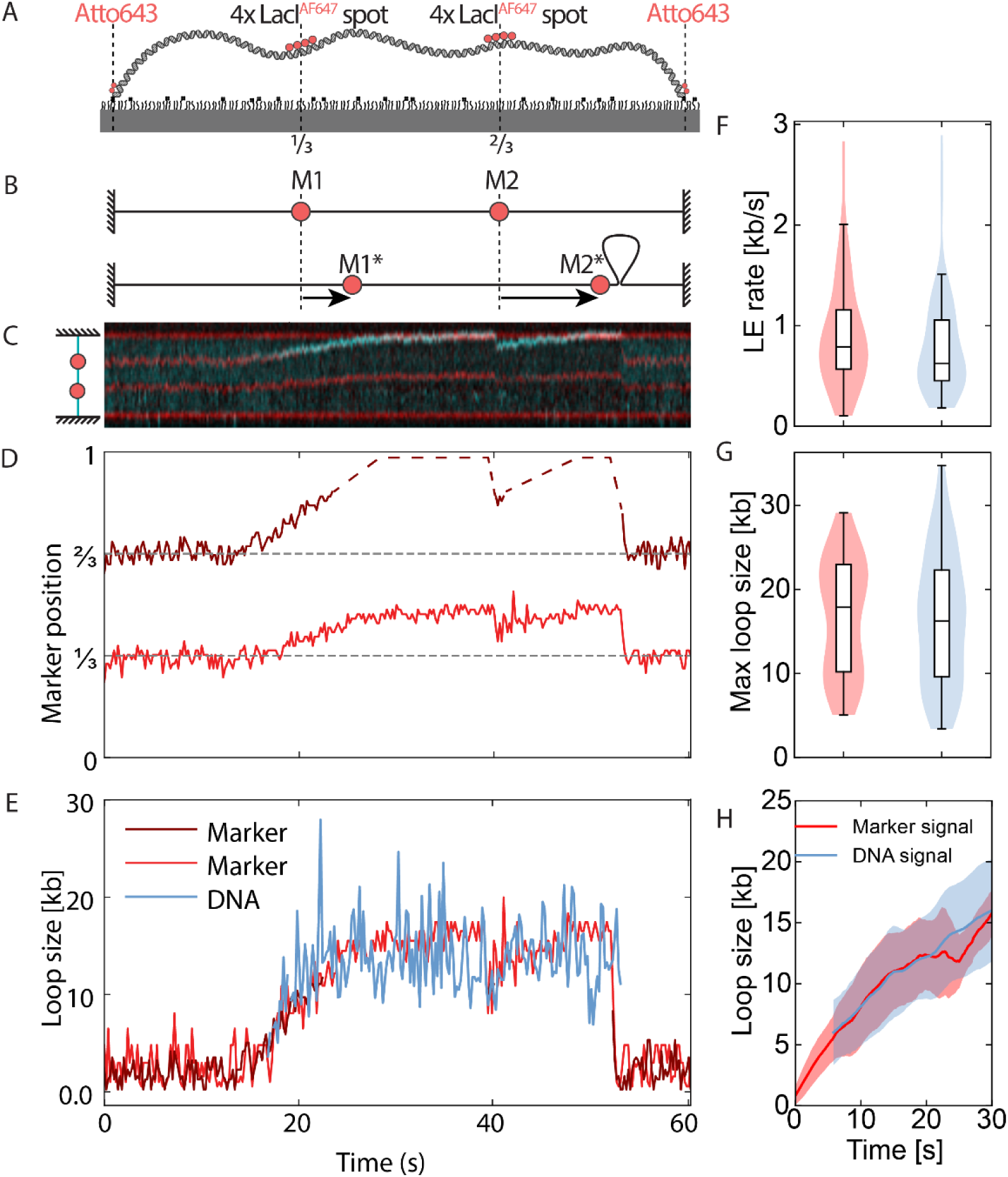
Modified loop extrusion assay based on spot label markers. **A)** Schematic representation of spot-labeled DNA constructs bound to the PEGylated glass surface. The 50 kb construct has Atto643-labeled biotin handles at either end for binding to streptavidin-coated surface. Two marker positions located at 1/3 and 2/3 along the construct contain four LacO sites each. The Atto643-labeled ends serve as static markers of the boundary of DNA, whereas the LacO sites serve as movable markers when bound by Alexa Fluor 647-labeled LacI. **B)** Schematic representation of the effect of an extruded loop on the movable markers. As a loop effectively removes a stretch of DNA from the contour of the tethered construct, the markers reposition and their positional shift is directly related to the amount of DNA in the loop. **C-E)** Example of loop extrusion on the spot-labeled constructs with 10 nM SYBR Gold. **C)** Kymographs of DNA/SYBR Gold signal (cyan), and the markers (red) that are pulled in towards the loop. Increasing cyan intensity is indicative of an extruded loop. **D)** Tracked marker positions and their shift relative to their original positions as indicated by the dashed gray lines. The position of one marker could not be tracked accurately due to close proximity to the boundary marker and its signal is replaced by the dashed line as a guide to the eye. **E)** Extracted loop size from the DNA signal and both markers. **F-H)** Loop extrusion rate, maximum loop size, and loop extrusion time calculated from both the marker signal and DNA signal in the 10 nM SYBR Gold dataset, which includes N=36 loop extrusion traces. None of the distributions showed any significant differences (Kolmogorov-Smirnov test p > 0.05).

where *M*^∗^ is the new marker position after movement. In our experimental setup,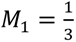 and 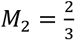 and their positional shifts relative to their original position is simply inferred from microscopy.

For both methods of obtaining the loop size, the loop size was analysed over time. The initial extrusion rate was calculated using the method that we here refer to as the “falling stick algorithm”. In this method, the obtained loop sizes are filtered with a Savitzky-Golay filter (window=11 frames, polynomial order=1) to reduce noise. Then, the loop extrusion rate was simply calculated as the maximum slope between the initial timepoint where loop extrusion started and subsequent timepoints. To minimize the effects of initial noise, the calculation was performed over at least the first 10 frames.

## Results

### Intercalator binding to DNA

In order to quantitatively compare different intercalating dyes and how their binding affects DNA’s mechanical properties, we start by determining their binding density on DNA in the typical buffer composition used in the loop extrusion assays (50 mM KGlu, 2.5 mM MgCl_2_, 1 mM TCEP, 0.25 mg/ml BSA, 20 mM Tris-HCl pH 7.5). As previously reported^8^, the concentration of salts in a buffer strongly impacts the binding affinity of intercalators. Intercalator binding to DNA follows the McGhee-von Hippel binding model:^47^

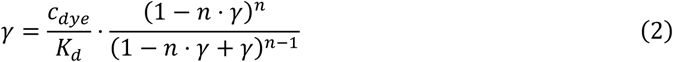

where *γ* is the fractional binding density (i.e. the number of intercalators per basepair), *c_dye_* the concentration of intercalator dyes in solution, *K_d_* the dissociation constant, and *n* the binding-site size, i.e. the dye’s footprint measured in basepairs. While titrating the dye at a constant DNA concentration in the loop-extrusion buffer, we measured the amount of binding from the fluorescence signal and fitted the dissociation constants from the curves, while fixing the binding site size to previously determined values (see Table 1 for details).

**Table 1.** Overview properties of the panel of studied dyes.

| **Intercalating dye** | **Abs. maximum [nm]** | **Em. maximum [nm]** | **Binding site size n** | **Affinity in LE buffer** | **Affinity in PBS** |
| --- | --- | --- | --- | --- | --- |
| SYBR Gold | 495 | 537 | 1.6 | 55 ± 16 nM | 206 ± 48 nM |
| YOYO-1 | 491 | 509 | 3.2 | 3 ± 5 nM | 4 ± 5 nM |
| SYTOX Orange | 547 | 570 | 2 | 149 ± 31 nM | 192 ± 30 nM |
| SYTOX Green | 504 | 523 | 3.2 | 17 ± 6 nM | 27 ± 4 nM |

We performed our measurements on a panel of four intercalating dyes that are commonly used and that show distinct spectral and chemical properties: SYBR Gold (SbGo), YOYO-1, SYTOX Orange (SxO), and SYTOX Green (SxG). From the measurements on these intercalators (Fig. 1B), we thus obtained a dissociation constant *K_d_* ranging from 3 to 150 nM. Specifically, we obtained a *K_d_* of 55±16 nM (mean ± standard deviation) for SbGo, 3.4±5 nM for YOYO-1, 149±31 nM for SxO, and 17±6 nM for SxG. To verify our method, we repeated our measurements in PBS for comparison with other previous reports. For SbGo, we then found a value of 206±48 nM, in good agreement with the previously reported 139-273 nM^9^; for YOYO-1 in PBS, we obtained 3.7±4.6 nM, in good agreement with the low nanomolar range previously reported^48^; and for SxO and SxG, we obtained 192±30 nM, and 27±4 nM, respectively – values that are also well in the range of a report using optical tweezers and slightly different monovalent salt concentrations^8^.

### Measurement of the persistence length of intercalated DNA at low tension

Next, we wondered whether it is possible to measure a change in the persistence length of intercalated DNA at very low tensions. Measuring the persistence length specifically in the low tension regime (≪ 1 pN) is relevant for the study of SMC motor proteins, whereas most methods that infer the persistence length involve stretching the polymer at high forces (5-60 pN), which alters the binding properties of these dyes^8,14,20^. To this end, we applied an approach to DNA that was previously established to analyse the persistence length of actin filaments and microtubules^49–52^. This approach makes use of imaging the bending fluctuations of semiflexible polymers, for which the persistence length and contour length are at the same length scale. For DNA with a persistence length of ∼50 nm, the original implementation of this approach is therefore only accessible to high-resolution techniques such as AFM on molecules in the same length scale^11^. However, we demonstrate that at the resolution provided by typical fluorescence microscopy, this framework can be expanded to the case of long flexible polymers such as the 50 kb DNA molecules in this work (see methods and supporting note 1 for details).

The approach involves to, first, trace the contour of the polymer and subsequently decompose it in half-harmonic bending modes. The variance of the coefficients of each of these modes is then related to the tension and persistence length of the polymer, based on^49^

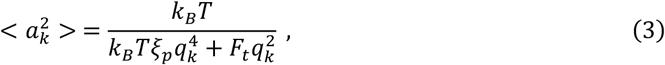

where the values *a_k_* are the bending mode coefficients for each mode *k*, *ξ_p_* is the persistence length, *F_t_* is the tension force, and *q_k_* the wavenumber for a mode *k*. This approach has been successfully applied to protein filaments^49–52^ and very short DNA molecules^11^ with a contour length that is roughly similar to the persistence length. However, it is unknown whether this approach can be applied to a polymer like >10 kb-long DNA, for which the persistence length is several hundred times smaller than the contour length.

To measure the persistence length of intercalated DNA using this approach, we applied an *in vitro* single-molecule microscopy technique that was previously established in our lab to study the supercoil^23^ and loop-extrusion dynamics^24^ of DNA, which makes use of intercalating dyes to visualize DNA. To this end, 50 kb DNA molecules were prepared with biotinylated ends for surface tethering using stable biotin-streptavidin interactions (Fig. 1C). These molecules were introduced into a flow channel under a mild flow, which stretches them to 30 ± 11% (median ± standard deviation) of the contour length at sub-piconewton forces (Fig. S2). These single DNA molecules were then imaged with a HILO fluorescence microscope. Recordings of tens of well-separated DNA molecules were made in the absence of any flow. Subsequently, the DNA tethers were traced and analysed over time to calculate their persistence length (Fig. S2). The molecules displayed clear fluctuations on sub-second timescales (Fig. 1D, top). Traces of the DNA molecules were decomposed into half-harmonic modes (Fig. 1D, bottom), yielding the coefficients for each bending mode. Then by simultaneously fitting the obtained coefficients for each recorded frame of each of the tens of single DNA molecules to equation 3, we obtained the individual tensions for each DNA molecule and a single persistence length per measurement condition. See Methods and supplemental note 1 for further details.

This analysis yielded persistence lengths as displayed in Fig. 1E. We find that the persistence length consistently increases for all dyes at increasing binding densities, numerically yielding higher values up to a 20% increase for some intercalators at the highest densities of 25%. In other words, we find, at very low DNA tension, that intercalating dyes stiffen the DNA upon binding. Comparing the various dyes, we measured the smallest increase of the persistence length, from 46 nm to 49 nm, for SYTOX Orange over the whole range of intercalator densities of 5-25% (i.e. up to 1 intercalator for every 4 base pairs), while YOYO-1 showed the largest increase to 57 nm. SYTOX Green showed a very similar effect on the persistence length as YOYO-1, while SYBR Gold distinctly showed an increase in persistence length already occurring at much lower binding densities (5-10%) than other dyes. These values are in good agreement with previous studies. Overall for low intercalator binding densities, we find values close to the reported 46 nm persistence length in the absence of any intercalator in a similar low-ionic buffer, as measured with magnetic tweezers^37^. For SYBR Gold, we found that the persistence length finally tended towards 55 nm, in accordance with the maximum persistence length of 57 nm found by a previous study^9^. The small increase in persistence length for SYTOX Orange is in line with earlier reports^8,42^, while SYTOX Green’s high similarity to YOYO-1 is remarkably distinct from recent findings using magnetic tweezers^42^.

### Measurement of the charge of intercalated DNA

The second biophysical property of DNA that is strongly impacted by intercalator binding is the net charge of the DNA molecule. As most DNA-binding proteins have a positively charged binding interface that interacts with the negative charge of DNA, we hypothesized that a net reduction of the charge upon intercalation may inhibit DNA binding – motivating a study of how intercalators modify the effective charge of DNA. Both the YOYO-1 and SYBR Gold dyes have a known positive charge, of +4 and +2, respectively^53,9^. However, both SYTOX dyes (SxO and SxG) are commercial products that had no reported value for their charge until recently, when values of +2 and +3 were reported respectively^42^. We measured the zeta potential of DNA molecules in the presence or absence of different dyes. The zeta potential offers a measure for the global surface charge of a charged particle in solution, which is calculated based on its electrophoretic mobility.

For bare DNA, we found the strongest zeta potential of -42.6 ± 1.6 mV (Fig. 1F). We next added intercalators at a fixed dye:bp ratio of 1:4 (except where noted). Addition of intercalating dye was found to systematically reduce the magnitude of the zeta potential in all cases, consistent with partial neutralization of the DNA backbone charge upon dye binding. However, the dyes did not shift the zeta potential in proportion to their molecular charges. SYBR Gold and SYTOX Orange (molecular charge z=+2) produced comparable shifts, to -38.7 ± 1.9 mV and -39.0 ± 1.4 mV, respectively, consistent with a 12.5% reduction of the surface charge density (Fig. S3). SYTOX Green (z=+3) and YOYO-1 (z=+4), instead clustered near a common larger shift of about -32 mV (-31.6±2.4 mV and -32.2±1.3 mV, respectively), consistent with a 25% reduction of the surface charge density (Fig. S3). Notably, this places SYTOX Green’s effective contribution at the +4 reference level, despite its reported molecular charge of +3. As a control, we also halved the YOYO-1 concentration (to 1:8 dye:bp), which produced an intermediate shift to -36.3± 0.4 approaching the +2e reference level in this comparison, as expected.

Interestingly, the effective charge neutralization in these experiments was smaller than expected based on the dye:bp ratio and the molecular dye charges, indicating that not all of a dye’s molecular charge contributes equally to neutralizing the DNA backbone charge. Nevertheless, it is clear that intercalation reduces the magnitude of DNA’s negative zeta potential, with the extent of the partial neutralization depending on the molecular charge but also on the specific dye. Specifically, SYTOX Green (Z=+3) was expected to result in an intermediate reduction in the zeta potential but instead it followed YOYO-1 closely. A likely explanation lies in the spatial distribution of positive charges within each intercalator. Structural data for these intercalators^9,42,53^ show that the positive charges are distributed across two distinct environments: the planar, base-stacked aromatic chromophore that is buried between adjacent base pairs upon intercalation, and the flexible side chains that project into the surrounding solvent. All dyes carry a single charge on their planar intercalated core (where bis-intercalating YOYO-1 has two such intercalated cores each with a single positive charge). SYTOX Orange and SYBR Gold each carry a single charge in their solvent exposed side chain, while SYTOX Green and YOYO-1 carry two positive charges in that region. We propose that these two arrangements are not electrokinetically equivalent, and yield different mobilities in the zeta potential measurements which probe surface charge. This hypothesis explains why SYBR Gold and SYTOX Orange (+1 charge on their sidechain) yield a 12.5% zeta potential reduction of the surface charge density, while YOYO-1 and SYTOX Green (+2 charge on their sidechain) both yield a 25% reduction.

### Condensin binding and DNA loop extrusion are reduced by intercalation

Having determined the binding affinity to DNA and the effects on the persistence length and DNA charge for our panel of dyes, we now turn to the intercalator-binding effects on the enzymatic activity of DNA-binding proteins on DNA. Specifically, we investigate their effects on condensin-mediated loop extrusion, which is a highly dynamic process in which repeated cycles of DNA binding, large conformational changes, and ATP hydrolysis within the condensin complex result in a stepwise increase in DNA loop size. To study loop extrusion, we employ our intercalator-based single-molecule visualization assay in which DNA is doubly tethered to a glass coverslip through biotin-streptavidin interactions and mild flow^24^. Condensin is then introduced into the channel together with ATP. Extruded loops are visible as bright puncta of the intercalator fluorescence signal, which represent locally more abundant regions of DNA. Kymograph analysis then allows the calculation of metrics such as the loop size, loop extrusion rate, loop extrusion time, and the tension on DNA. At a fixed intermediate intercalator density of 10%, loops were observed for all intercalating dyes (Fig. 2A), demonstrating that intercalation itself is compatible with DNA binding and enzyme activity of condensin. Indeed, previous studies already demonstrated loop extrusion with the intercalating dyes SYTOX Orange^24^, SYTOX Green^37^, and YOYO-1^54^.

From single-molecule kymographs, we measured the initial loop-extrusion rate, the maximum loop size (i.e. the loop size upon saturation of the loop-extrusion process), and total loop extrusion time (defined as the time between the start of loop extrusion and reaching the plateau in loop size) on DNA molecules that were stretched by ∼30-50% of their contour length (Fig. 2B-D). In this tension regime, we found median loop-extrusion rates ranging from 0.5 to 1 kb/s, with values for SYTOX Orange at the higher end of this range and SYTOX Green at the lower end. We did not measure any significant differences between the loop extrusion rates, maximum loop size or total loop extrusion time (Kolmogorov-Smirnov p > 0.05 for any pair). The maximum loop size was found to be similar for all dyes, with a value broadly distributed around 15 kb. The total loop extrusion times were similarly broadly distributed with a value ranging in the tens of seconds. While the loop extrusion times varied significantly across the different dyes overall (Kruskal–Wallis, p = 0.024), no significant differences were identified between any specific pair of dyes (pairwise Kolmogorov–Smirnov test). Note that, for consistency with previously reported values in the literature, we above reported the loop extrusion rate and loop sizes in base pairs, even though intercalator binding lengthens the DNA tether, increasing the total contour of intercalated DNA. Supplemental figure S4 reports our findings instead in contour length, i.e. the length of the DNA basepairs and the additional intercalated dyes together. Overall, we could not identify any differences in loop-extrusion properties between the dyes at a fixed intermediate binding density of 10%.

To test the overall enzymatic activity of condensin, we measured the bulk ATP hydrolysis rate of DNA-condensin mixtures in the presence of intercalators. We found that the ATPase rate negatively correlates with intercalator binding (Fig. 2E, Fig. S5). Remarkably, two groups of intercalators showed a distinct dependence on the intercalator binding density: YOYO-1 and SYTOX Green showed a clearly stronger inhibition of the ATPase rate than SYBR Gold and SYTOX Orange. This correlates with our earlier observation that YOYO-1 and SYTOX Green have a stronger effect on the zeta potential of DNA as opposed to SYBR Gold and SYTOX Orange. Indeed, when we consider the net charge neutralization due to intercalation rather than binding density alone, we find that all datapoints collapse on a single line (Fig. 2E inset). The data strongly support a role for the net charge of DNA in stimulating condensin’s ATPase activity, as dyes that neutralize more of the DNA charge suppress condensin’s ATPase activity more strongly.

Because the loop extrusion rate did not show major differences among the dyes while counterintuitively the ATPase activity did reveal two distinct groups of intercalators, we next tested whether the difference in ATPase activity stemmed from a difference in initial DNA binding activity. Indeed, we observed that intercalation negatively correlates with condensin binding to DNA, as measured both in an electrophoretic mobility shift assay and a mass photometry assay (Fig. 2F, Fig. S6-7). We observed that YOYO-1 and SYTOX Green inhibit condensin’s DNA binding more than SYBR Gold and SYTOX Orange do (Fig. 2G, Fig. S6), in good agreement with the ATPase activity and zeta potential measurements.

Reconciling the measured loop extrusion rates, ATPase activity, and DNA-binding activity, we conclude that intercalator binding competes with condensin binding to DNA and that, once condensin has bound to DNA, it has a set loop extrusion rate that is fairly independent of the DNA charge. Furthermore, we conclude that the bulk ATPase activity is set mostly by DNA binding rather than by loop extrusion. Finally, we note that in the standard loop-extrusion assay, the intercalator concentration is inherently coupled to signal quality (as higher densities improve the signal-to-noise ratio), which confounds any density-dependent trends and prevents a detailed study of a systematic variation of loop parameters versus intercalator density. Notably, the standard assay precludes the measurement in the complete absence of intercalator, leaving this baseline inaccessible. These limitations call for a fundamentally different approach to measuring loop extrusion dynamics *in vitro,* which we present next.

### A new loop-extrusion assay without intercalating dyes

We developed a new in vitro loop-extrusion assay that uses no dyes whatsoever. The standard assay requires intercalators to visualize the DNA and quantify loop extrusion properties. While loop extrusion was also observed (though not quantitatively studied) by labelling the DNA with fluorophores that are covalently attached to bases^38^, that method has its drawbacks because of the necessity for chemically modifying a large portion of the base pairs.

Here, we present a loop-extrusion assay where only a few local spots are fluorescently labelled (Fig. 3A). These spot labels act as markers that can be tracked while their positions change over time as a condensin extrudes a DNA loop somewhere else in the DNA molecule (Fig. 3B). The markers move over time because the tethered DNA tightens up as a loop is being extruded by the condensin at one spot. Their positional shift directly relates to the size of the extruded loop, so the loop-extrusion properties can be extracted from the positions of the markers alone in the absence of fluorescent signal from any intercalators (see Methods and Fig. S8 for an extensive overview of these calculations and examples).

Figure 3 illustrates the new assay. For testing purposes, we recorded condensin-induced loop-extrusion events with this assay in the presence of an intercalator (SYBR Gold at an intercalator density of 10%), and separately analysed the extruded loops based on the intercalator signal as well as based on the marker positions. In the example shown in Fig. 3A-E, a loop initiated close to one of the markers and the loop extruded towards the proximal DNA end. Both markers moved along with the loop and their positional shift corresponded directly to the size of the extruded loop (Fig. 3D-E). The markers clearly moved upon the initiation and continuation of loop extrusion. Strikingly, the marker detection provided a more sensitive measure of loop extrusion, as it appeared to exhibit a less noisy signal (compare the red and blue traces in Fig. 3E). Furthermore, the new assay typically detected loop extrusion many frames before any loop could be detected in the intercalator fluorescence signal (which detected loops only above a finite intensity threshold of a minimal (few kb) amount of DNA above background).

Importantly, the two methods yielded very similar estimates of the loop-extrusion rate and maximum loop size (Fig. 3F-H, Fig. S9). While these properties did not show any significant differences between the quantities obtained from the markers or DNA fluorescence, the loop extrusion rate was slightly lower when quantified by DNA fluorescence. This small difference is readily explained from the detection threshold for the DNA fluorescence signal, which allows tracking the loop only from a slightly later timepoint, when the loop extrusion rate has slowed down already a bit. To quantify this effect, we ensemble-averaged all trajectories from the same dataset (Fig. 3H and methods). In these ensemble averages, the loop size calculated from the marker displacement started at 0 kb loop size, while the loop sizes calculated from the intercalated DNA signal could only be resolved starting from ∼5 kb.

We thus find that this new loop extrusion assay – which uses spot label markers rather than intercalated DNA signal to probe loop extrusion and measure loop size – is an excellent and in fact more accurate method to track the initial phase of loop extrusion, which is independent of DNA intercalation. It furthermore turns intercalator density from a necessity in the assay into a free experimental parameter.

### DNA loop extrusion by SMCs is hampered by intercalator binding

Having validated the novel assay, we next applied it to measure loop extrusion in the absence of intercalator binding (Fig. 4A), providing an unobstructed view of loop extrusion dynamics that is by definition unaffected by the intercalator concentration.

**Figure 4.**
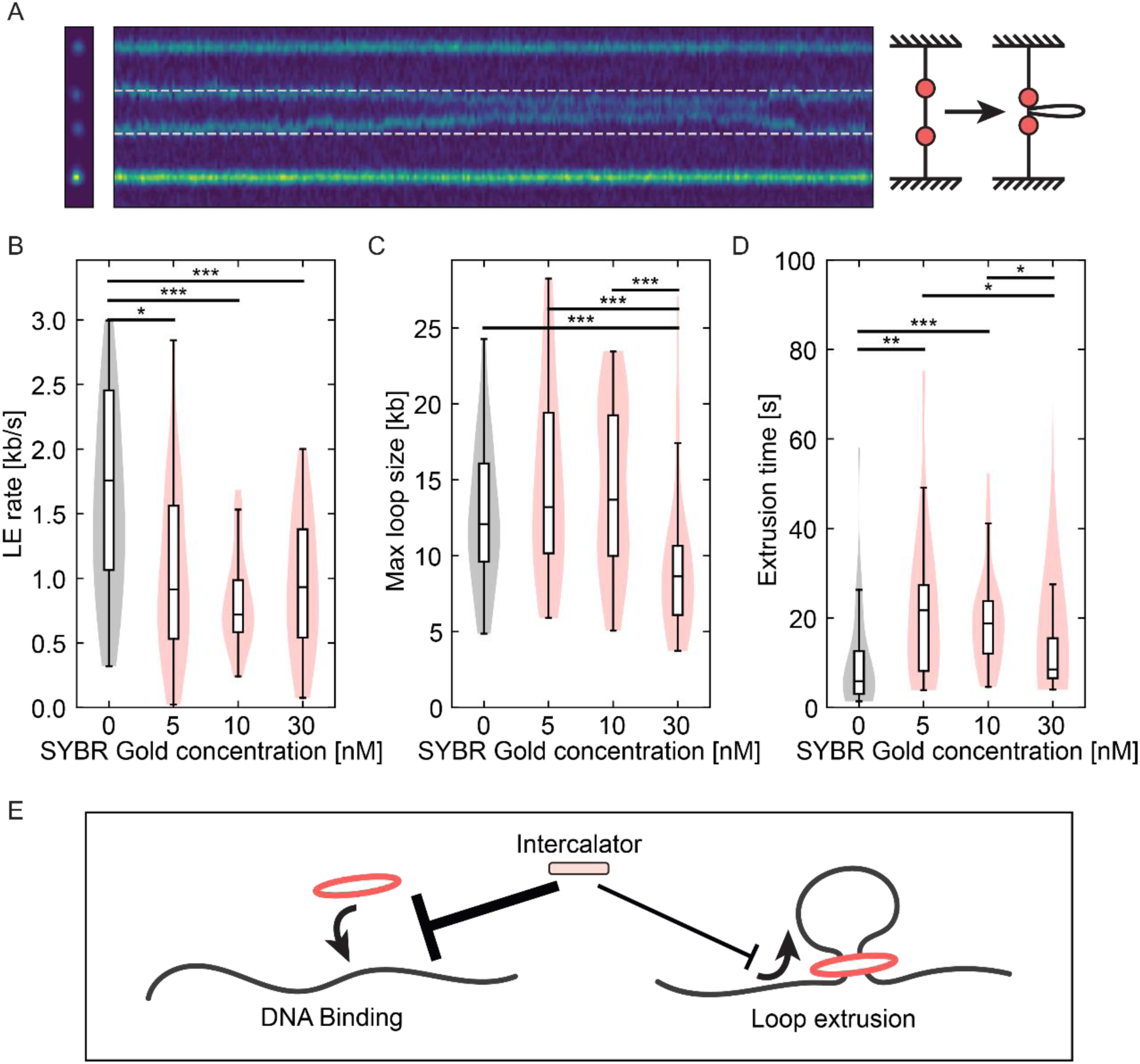
Loop extrusion in the absence of intercalating dyes. **A)** Example of markers moving in the presence of condensin. The kymograph represents 60s of recording and the markers both move towards the middle, indicating a loop being formed between them. **B-D)** Loop extrusion rate, maximum loop size, and loop extrusion time in the absence and presence of SYBR Gold at indicated concentrations. The indicated SYBR Gold concentrations correspond roughly to binding densities of 5%, 10%, and 25%, respectively. Significant differences (Kolmogorov-Smirnov) are indicated as follows: *: p <0.05, **: p<0.01, ***: p<0.001. N= 39, 25, 36, 27, respectively. **E)** Intercalator binding mostly impacts DNA binding, and affects the loop extrusion rate.

Strikingly, as shown in Fig. 4B, we observed a significantly faster loop extrusion rate in the absence of dye, namely a rate of 1.75 ± 0.26 kb/s (median ± SEM, mean was 2.16 kb/s) compared to median values of 0.7-0.9 kb/s in the presence of SYBR Gold at concentrations between 5 and 30 nM (corresponding to intercalator binding density of 5 to 25%). The SYBR Gold intercalators thus impede the loop extrusion rate by roughly a factor of 2. In contrast to the ATPase and DNA binding activities, we did not find a simple linear intercalator-density effect on the loop extrusion rate, but rather a large attenuation of the rate when it was measured in the presence of intercalating dyes. We did not observe a significantly different maximum loop size when comparing the various conditions with and without intercalator, with the exception of the highest intercalator concentration of 30 nM which showed a lower maximum loop size (Fig. 4C). The combination of varying loop extrusion rates and the same maximum loop sizes resulted in corresponding differences in the overall loop extrusion times (Fig. 4D).

In all, we concluded that the presence of intercalators hampers loop extrusion, as we find a twice larger loop extrusion rate in the absence of intercalator. As intercalator fluorescence detection was used in all previous *in vitro* estimates of the loop-extrusion rates, this new finding shows that the actual loop-extrusion rate is much higher than reported before *in vitro*.

## Discussion

We measured the effects of intercalating dyes on DNA and DNA-binding proteins. Specifically, we studied how intercalating dyes in DNA affect *S. cerevisiae* condensin, a highly dynamic molecular motor that extrudes loops of DNA. We quantified how four distinct intercalating dyes modify both the persistence length and net charge of DNA. We investigated how intercalation modifies condensin’s interactions with DNA and we quantified the effects on ATPase activity, DNA binding, and loop extrusion. To measure the effect of intercalators, we developed a new single-molecule assay with local markers that, unlike the standard *in vitro* single-molecule DNA-visualization assay^24–26^, does not depend on intercalator signal to measure loop extrusion properties. We found that this method more accurately tracks DNA loops from the onset of loop extrusion, where the intercalator signal is still indistinguishable from background, and we deduced a roughly twice as fast loop extrusion rate in the absence of intercalating dyes.

### Changes in DNA mechanics upon intercalation

We used a novel method to analyse the persistence length of DNA molecules based on examining the bending fluctuations of stretched DNA. Our analysis revealed that intercalation tends to stiffen the DNA. At low intercalator binding densities, the measured persistence length was consistent with earlier measurements in a similar buffer^37^, indicating that this method is an accurate method that agrees well with other techniques. The biggest advantage of tracking the lateral fluctuations of surface-tethered DNA molecules is that we can probe the intrinsic mechanical properties of DNA in the regime of the very low tensions that are applied on the molecules (a regime particularly relevant to SMC proteins). A typical 50 kb DNA tether in this assay may be stretched 6-9 μm, which corresponds to a tension of 50-100 fN (0.05-0.1 pN), much lower than the forces used in optical or magnetic tweezers experiments. While one downside of this method is the necessary visualization of DNA molecules, we demonstrate that we were able to image DNA molecules at sufficient signal-to-noise at only 5% intercalation of SYBR Gold or SYTOX Orange, and we extracted persistence lengths here that were close to those reported using other techniques.

Optical and magnetic tweezers assays (that typically operate in the 1-50 pN tension regime) reported different values for the persistence length in the presence of intercalators. A complication is that intercalator binding is itself force-dependent: for example, ethidium bromide was demonstrated to have a much smaller binding site and higher affinity at 50 pN compared to 5 pN, strongly altering key binding parameters as a function of tension^14^. As a result, the contour length changes along a force-extension curve and when such data are fitted with a single WLC of fixed contour length, this force-dependent lengthening is fit as due to an apparent flexibility, which can yield strongly reduced persistence lengths such as the 5.7 nm value reported for EtBr^14^. However, AFM studies of ethidium-intercalated DNA showed an increase rather than decrease of the persistence length^11^. Accordingly, in one magnetic tweezers study where the DNA tension was limited to a medium tension regime of <5 pN, a persistence length of ∼59 nm was found at saturating SYBR Gold concentration^9^. Similarly in our study, we found a limiting persistence length around 55 nm with SYBR Gold. While previous persistence length measurements of intercalated DNA using stretching techniques yielded variable values for different tension regimes, we consistently observed an increased stiffness upon intercalation, as the tension on DNA was held consistently low in our fluctuation analysis.

An important consequence for condensin-DNA interaction is that DNA-binding proteins encounter a charged DNA surface with an effective zeta potential that is the combined result of DNA charge, intercalator charge, and local counterions.

### SYBR Gold and SYTOX Orange are the least invasive intercalating dyes to study protein-DNA interactions

We demonstrated that *S. cerevisiae* condensin’s DNA binding, ATPase activity, and loop extrusion rate are strongly impacted when using higher intercalator binding densities. This raises important questions for their use to study protein-DNA interactions. Practically, it is important to consider which specific intercalator is most appropriate.

Overall, we find that SYBR Gold and SYTOX Orange had the smallest effect on the net charge of DNA. Of all dyes tested, SYTOX Orange had the smallest effect on DNA persistence length across all concentrations. SYBR Gold had a similarly small effect at low intercalation densities, but stiffened the DNA more at higher densities. Furthermore, YOYO-1 has a remarkably high residence time (48 seconds in a 0.1 M NaCl buffer) while the other dyes exchange much more rapidly^8^. Combining these observations, we conclude that SYBR Gold and SYTOX Orange are the most suitable intercalating dyes for studying protein-DNA interactions.

### The use of intercalating dyes in SMC loop-extrusion assays

While loop extrusion is qualitatively not affected by intercalating dyes, quantitative values for binding and looping rates of SMC-driven DNA loop extrusion differ from values that were previously reported in literature. For their use in the single-molecule loop extrusion assay, we find that SYTOX Orange and SYBR Gold are excellent options for visualization. Spectral considerations are relevant when imaging multiple proteins alongside the DNA. In order to limit the effects of bleed-through, DNA is typically imaged with the smallest wavelength available. Even though SYTOX Green has been used in the past for exactly this reason^37,40,41,55–58^, we find here that SYBR Gold is more appropriate due to its milder effects on DNA and SMC-mediated loop extrusion. While SYBR Gold and SYTOX Orange both have quantitatively similar effects on condensin and loop extrusion, SYBR Gold is brighter and allows imaging in an additional fluorescent wavelength and is thus the preferred dye. Notably, we analysed a finite set of 4 intercalating dyes, comprising the most commonly used ones, and hence there is always the possibility that yet other intercalators are more suited for single-molecule visualisation.

We find that SYBR Gold is the typically preferred intercalating dye, which can be used at minimal binding densities (5 nM, corresponding to ∼5% binding density). At this concentration, SYBR Gold results in sufficient signal for the analysis of loop extrusion, while the DNA persistence length and charge are not yet much affected. Other intercalators need to be used at much higher binding densities for adequate visualization, and their use for SMC dynamics studies should be limited as much as possible. A useful rule-of-thumb is to use the lowest possible intercalator concentration that still allows for the quantification of whichever process is studied.

We find that intercalators inhibit DNA loop extrusion by SMCs in two ways: through initial DNA binding of the SMC and through the subsequent loop-extrusion process (while the final loop sizes remain unaffected). It is clear that SMC complexes are challenged by intercalator binding to DNA, an important consideration that has to be taken into account in modelling and extrapolating single-molecule results to *in vivo* situations.

This raises the question of how fast SMC complexes extrude loops when they are not challenged by intercalators at all. Using a novel intercalator-free assay, we found a surprisingly high value for the loop extrusion rate of 1.75 kb/s, which is roughly double the value that was obtained with intercalators. Notably, that high value is well in line with mitotic loop extrusion rates measured *in vivo*, which are in the range of 1-3 kb/s^33,59^. While there is no *in vitro* measurement of the loop extrusion rate on chromatinized DNA, we surmise that loop extrusion would exhibit a similarly sped up loop-extrusion velocity since nucleosomes do not block loop extrusion^38,39^.

Our finding challenges previous quantifications of the loop-extrusion rate. In Table 2, we compiled a comprehensive list of all *in vitro* single-molecule studies of loop extrusion, alongside the reported loop extrusion rates and an estimate of intercalator binding density. It can be seen that SYTOX Orange has been used most frequently in these assays. The typical concentrations for loop-extrusion visualisation correspond mostly to relatively high intercalator densities, which have strong effects on the DNA binding, ATPase, and loop-extrusion activities. For example, we estimate that previous single-molecule loop extrusion experiments were performed with SYTOX Orange binding densities in the higher range of 20-35%. Previously reported values are in line with our current measurements at similar intercalator concentrations. Note, however, that this comparison does not take into account the effect of initial DNA tension, which is an additional key factor that can strongly alter the loop-extrusion rate.

**Table 2.** Overview of single-molecule loop extrusion studies with intercalators.

| SMC complex | Dye | Conc. [nM] | Salts | Binding density | LE rate [kb/s] | Ref. |
| --- | --- | --- | --- | --- | --- | --- |
| *S. cerevisiae* Condensin | None | N/A | 20 mM Tris-HCl, 50 mM KGlu | N/A | 1.75 | This work |
| *S. cerevisiae* Condensin | SYBR Gold | 10 | 20 mM Tris-HCl, 50 mM KGlu | 10% | 0.72 | This work |
| *S. cerevisiae* Condensin | SYTOX Orange | 30 | 20 mM Tris-HCl, 50 mM KGlu | 10% | 1.02 | This work |
| *S. cerevisiae* Condensin | SYTOX Green | 5.25 | 20 mM Tris-HCl, 50 mM KGlu | 10% | 0.56 | This work |
| *S. cerevisiae* Condensin | YOYO-1 | 1.1 | 20 mM Tris-HCl, 50 mM KGlu | 10% | 0.94 | This work |
| *S. cerevisiae* Condensin | SYTOX Orange | 500 | 50 mM Tris-HCl, 50 mM NaCl | 35.6%<sup>a</sup> | 0.6 | Ganji 2018<sup>24</sup> |
| Human Cohesin | SYTOX Orange | 250 | 50 mM Tris-HCl, 50 mM NaCl | 30.5%<sup>a</sup> | 0.8-1 | Davidson 2019<sup>25</sup> |
| Human Cohesin | SYTOX Orange | 220 | 50 mM Tris-HCl, 50 mM NaCl | 29.4%<sup>a</sup> | 1 | Davidson 2023<sup>25</sup> |
| *S. cerevisiae* SMC5/6 | SYTOX Orange | 100 | 40 mM Tris-HCl, 50 mM NaCl, 2.5 mM MgCl<sub>2</sub> | 22.2%<sup>b</sup> | 0.4 | Barth 2025<sup>40</sup> |
| *S. cerevisiae* SMC5/6 purified from *E. coli* | SYTOX Orange | 100 | 40 mM Tris, 100 mM NaCl, 7.5 mM MgCl<sub>2</sub> | 21.5%<sup>c</sup> | 0.7 | Barth 2025<sup>40</sup> |
| *S. cerevisiae* SMC5/6 | SYTOX Orange | 200 | 40 mM Tris-HCl, 100 mM NaCl, 7.5 mM MgCl<sub>2</sub> | 28%<sup>c</sup> | 1.3 (SMC5/6 dimer) | Pradhan 2023<sup>26</sup> |
| Human Cohesin | YOYO-1 | 0.2-0.5 | 40 mM Tris-HCl, 50 mM NaCl | 4.5-8.5%<sup>d</sup> | 0.5* | Kim 2019<sup>54</sup> |
| Human Condensin I | YOYO-1 | 0.5 | 40 mM Tris-HCl, 125 mM NaCl, 5 mM MgCl<sub>2</sub> | 8%<sup>e</sup> | 0.9* | Kong 2020<sup>60</sup> |
| Human Condensin II | YOYO-1 | 0.5 | 40 mM Tris-HCl, 125 mM NaCl, 5 mM MgCl<sub>2</sub> | 8%<sup>e</sup> | 0.2* | Kong 2020<sup>60</sup> |
| *P. aeruginosa* JetABC | SYTOX Orange | 100 | 40 mM Tris-HCl, 100 mM NaCl, 7.5 mM MgCl<sub>2</sub> | 21.5%<sup>c</sup> | 0.5 (dimer) | Pradhan 2025<sup>61</sup> |
| *P. aeruginosa* JetABCD | SYTOX Orange | 100 | 40 mM Tris-HCl, 100 mM NaCl, 7.5 mM MgCl<sub>2</sub> | 21.5%<sup>c</sup> | 0.7 (dimer) | Pradhan 2025<sup>61</sup> |

\* These values were obtained using a different technique (DNA curtains) which imposes a non-uniform DNA tension and constant buffer flow, which will affect the results.
<sup>a</sup> The affinity to DNA was determined experimentally and McGhee-von Hippel (MvH) parameters were: $n=2$ , $K_d=179$ nM.
<sup>b</sup> The affinity to DNA was approximated based on the buffer marked with (<sup>a</sup>), MvH parameters were: $n=2$ , $K_d=179$ nM.
<sup>c</sup> The affinity to DNA was approximated based on PBS buffer, MvH parameters were: $n=2$ , $K_d=192$ nM.
<sup>d</sup> The affinity to DNA was approximated based on the loop extrusion buffer at 20 mM Tris-HCl and 50 mM KGlu, MvH parameters were: $n=3.2$ , $K_d=3.39$ nM.
<sup>e</sup> The affinity to DNA was approximated based on PBS buffer, MvH parameters were: $n=3.2$ , $K_d=3.7$ nM

While we examined only one SMC complex, *S. cerevisiae* condensin, in our current study, we expect that our findings are generally applicable to all SMC complexes and other DNA motor proteins.

It is important to note that SMC complexes are a unique class of motor proteins that make relatively sparse contacts with DNA while stepping along DNA in very large steps (hundreds of bp/step), compared to e.g. helicases or polymerases that contact every basepair that they step over. For example, a previous study on the effect of intercalator binding on DNA polymerase activity found that the intercalator’s off-rate had a strong effect on enzyme activity^8^ as intercalator binding blocked the polymerase, which could only continue once the intercalator unbound. Our data, by contrast, indicate that SMC binding to DNA and loop extrusion are mostly impacted by the overall charge of intercalated DNA rather than by the intercalator off-rate. This supports a model in which SMC complexes take large steps along DNA driven largely by nonspecific electrostatic contacts irrespective of individual intercalators as physical roadblocks.

Altogether, our results show that intercalating dyes are not merely passive reporters in single-molecule assays because they influence the DNA mechanics. Encouragingly, the qualitative picture of SMC-driven loop extrusion that has emerged from intercalator-based assays remains intact, but quantitatively, we find that condensin extrudes DNA about twice as fast as previously thought once freed from intercalating dyes. With this loop extrusion rate as a quantitative benchmark and our practical guidelines for dye choice and concentration, we anticipate that this work will allow future single-molecule studies to capture the intrinsic behaviour of SMC complexes.

## Data availability

Data and code generated for this publication are deposited at the 4TU data repository and will be made available publicly at publication.

## Acknowledgments

We thank Ashmiani van den Berg for purifying S. cerevisiae condensin; Eli van der Sluis for purifying fluorescently labelled LacI dimers; Thejas Satheesh, Theo Koenig, Lisa Warners, Marjon van Ruiten, Allard Katan, and Beatriz Orozco Monroy for discussions; and Veerle and Tilly Schaap for support. We acknowledge the use of LLMs (Claude, Anthropic) for brainstorming, editing, and code improvement. We also acknowledge the NWO-XL project ‘Resolving the fundamental building principles of the genome’ and the ERC Synergy project GeneMotors for funding.

## Author Contributions

BTA: Conceptualization, methodology, investigation, formal analysis, writing. JvdT: Resources, methodology, writing – review & editing. CD: Supervision, funding acquisition, writing.

