## Supplementary information for "Intercalator effects on DNA loop extrusion by SMC proteins"

### Supporting Information Text

#### Supporting note 1: Derivation of fitting equations for lateral fluctuation analysis of DNA

While the lateral fluctuation analysis as introduced in the main text was derived in the context of semiflexible polymers[1–4], it is possible to extend the analysis to the case of more flexible polymers under specific conditions. Semiflexible polymers are those polymers for which the contour length $L$ is of the same order as the persistence length $\xi_p$. Notably, the long 50 kb DNA molecules in this study have a $L/\xi_p$ ratio of 340, placing them well in the flexible regime. However, when the polymer is analysed at sufficient coarse-graining, the lateral fluctuation bending-mode decomposition is still applicable for these molecules where $L \gg \xi_p$. In the derivation of the fitting equations below, we take extra care to stipulate the origin of this condition.

The Hamiltonian for a weakly bent polymer is defined as follows [2]:

$$\mathcal{H} = \frac{1}{2}\int_0^L (k_B T\xi_p(\nabla^2 y)^2 + F_t(\nabla y)^2)\, dx\,, \tag{1}$$

where $y = y(x,t)$ are the lateral fluctuations, $k_B$ Boltzmann's constant, $T$ the absolute temperature, and $F_T$ is the tension force. Following Gittes and colleagues[1], we perform a Fourier-mode decomposition (here choosing a sine basis function rather than cosine), yielding a summation over the different harmonic degrees of freedom:

$$\phi_k = \sqrt{\frac{2}{L}}\sin(q_k x)\,, \tag{2}$$

$$y(x,t) = \sum_k a_k\, \phi_k\,, \tag{3}$$

where $q_k = \frac{k\pi}{L}$ is the wavenumber, the prefactor $\sqrt{\frac{2}{L}}$ was chosen for normalization, and the bending mode coefficient $a_k$ represent the relative contribution of each bending mode. This decomposition allows us to write the Hamiltonian in eq. 1 in terms of the basis functions:

$$\mathcal{H} = \sum_k \frac{1}{2}\int_0^L k_B T\xi_p(\nabla^2 a_k\phi_k)^2 + F_t(\nabla a_k\phi_k)^2 dx, \tag{4}$$

$$= \sum_k \frac{1}{2}(k_B T\xi_p\, q_k^4 + F_t q_k^2)\, a_k^2. \tag{5}$$

Next, we use the equipartition theorem (which states that each quadratic mode, i.e. degree of freedom, contributes $\frac{1}{2}k_B T$ to the Hamiltonian) to arrive at our main fitting equation:

$$\boxed{< a_k^2 >= \frac{k_B T}{k_B T\xi_p q_k^4 + F_t q_k^2}}\,. \tag{6}$$

So, by tracing DNA molecules over time to obtain the values $y(x,t)$, decomposing their bending modes using eqs. 2 and 3, we obtain the coefficients $a_k$. Then from these coefficients and the known worm-like chain tension at a given extension[5], we can obtain the persistence length $\xi_p$ from eq. 6 from the first bending mode alone. The only limitation on validity in this derivation is made in the definition of the Hamiltonian in eq. 1 (weakly bent polymer). While this indeed is not generally applicable, we note that the first bending mode is already sufficient to obtain the persistence length. Indeed, the first bending mode represents a half-harmonic, which in the case of a 17 μm DNA molecule with a persistence length of roughly 50 nm is by definition a weak bend at this level of coarse graining.

Put formally, the mode wavenumber $q_k = \frac{k\pi}{L}$ defines the length scale of the fluctuations in mode $k$. As such, as long as the condition $\xi_p q_k \ll 1$ is fulfilled, this analysis holds (which is nearly equivalent to the statement that $L \gg \xi_p$). From this relationship, we find that in fact several lower orders may be used for this analysis. However as discussed in the next subsection, practical constraints limit the number of modes that can be used.

A second parameter that can be extracted from the mode decomposition is the time decay of fluctuations. Following the derivation of Kroy and Frey[4], we write the overdamped Langevin equation for lateral motions as:

$$\zeta_\perp \frac{\partial y(x,t)}{\partial t} = -\frac{\delta \mathcal{H}}{\delta y(x,t)} + \mathcal{N}(x,t), \tag{7}$$

where the left-hand term is the damping due to viscous drag, the first right-hand term is the functional derivative of the Hamiltonian with respect to the lateral displacement, and the last term is the Gaussian thermal noise. Working out the functional derivative and decomposing into harmonic modes, we then write:

$$\zeta_\perp \frac{\partial y(x,t)}{\partial t} = -k_B T \xi_p \frac{\partial^4 y(x,t)}{\partial x^4} - F_t \frac{\partial^2 y(x,t)}{\partial x^2} + \mathcal{N}(x,t), \tag{8}$$

$$\zeta_\perp \frac{da_k}{dt} = -\underbrace{\left(k_B T \xi_p\, q_k^4 + F_t\, q_k^2\right)}_{\equiv \frac{\zeta_\perp}{\tau_k}} a_k + \mathcal{N}(x,t). \tag{9}$$

So we find that the mode coefficients $a_k$ for each molecule have an autocorrelation function with a mode-specific time constant

$$\tau_k = \frac{\zeta_\perp}{k_B T \xi_p\, q_k^4 + F_t\, q_k^2}. \tag{10}$$

In principle, this relationship could also be used to measure the persistence length. Determining the drag coefficient $\zeta_\perp$ is nontrivial, however, since it varies depending on the relative stretching of the DNA molecule due to increased proximity to the surface at higher stretching. Hence, we cannot use the above relation in our fitting routine directly. However, we may still use the autocorrelation of mode coefficients as a quality control metric.

Moreover, the relation derived above sets a direct practical limitation on the measurements. The exposure time of the camera must be such that a traced DNA molecule can be regarded as static. If the exposure time is set too high, motion blurring starts to occur. For a DNA molecule stretched to 50% of its contour length and a canonical persistence length of 50nm, we estimate that $\tau_1 =$ 315 ms and $\tau_2 = 79$ ms. While the first bending mode can be accurately captured at an exposure time of 100 ms, we see that the second bending mode already decays within that single frame.

**Supporting Figures**

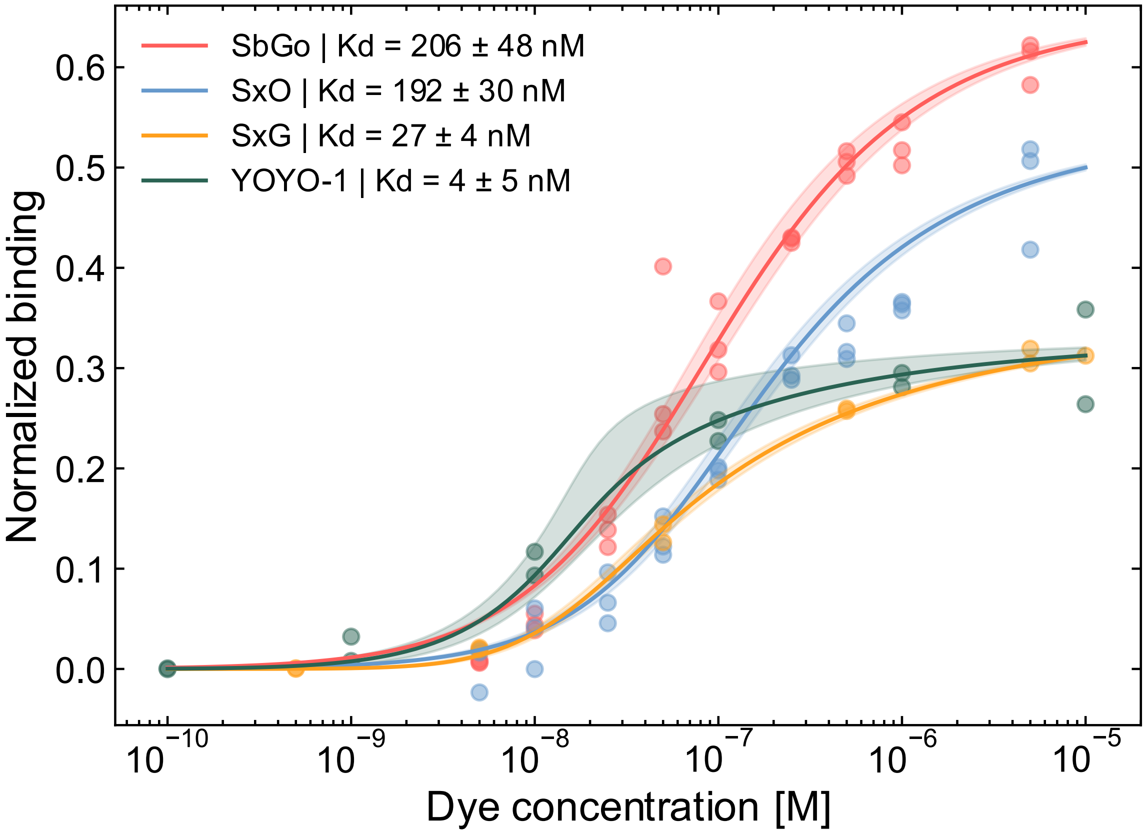


**Fig. S1. Intercalator binding in PBS.** Titration curves for SBYR Gold (SbGo), SYTOX Orange (SxO), SYTOX Green (SxG), and YOYO-1 dyes in PBS (phosphate buffered saline). Fitted affinities from the McGhee-von Hippel are given in the legend with the *Kd* as fit ± standard deviation. Individual measurements are shown as dots, fitted curves are shown as lines, and standard deviation is indicated as the shaded area.

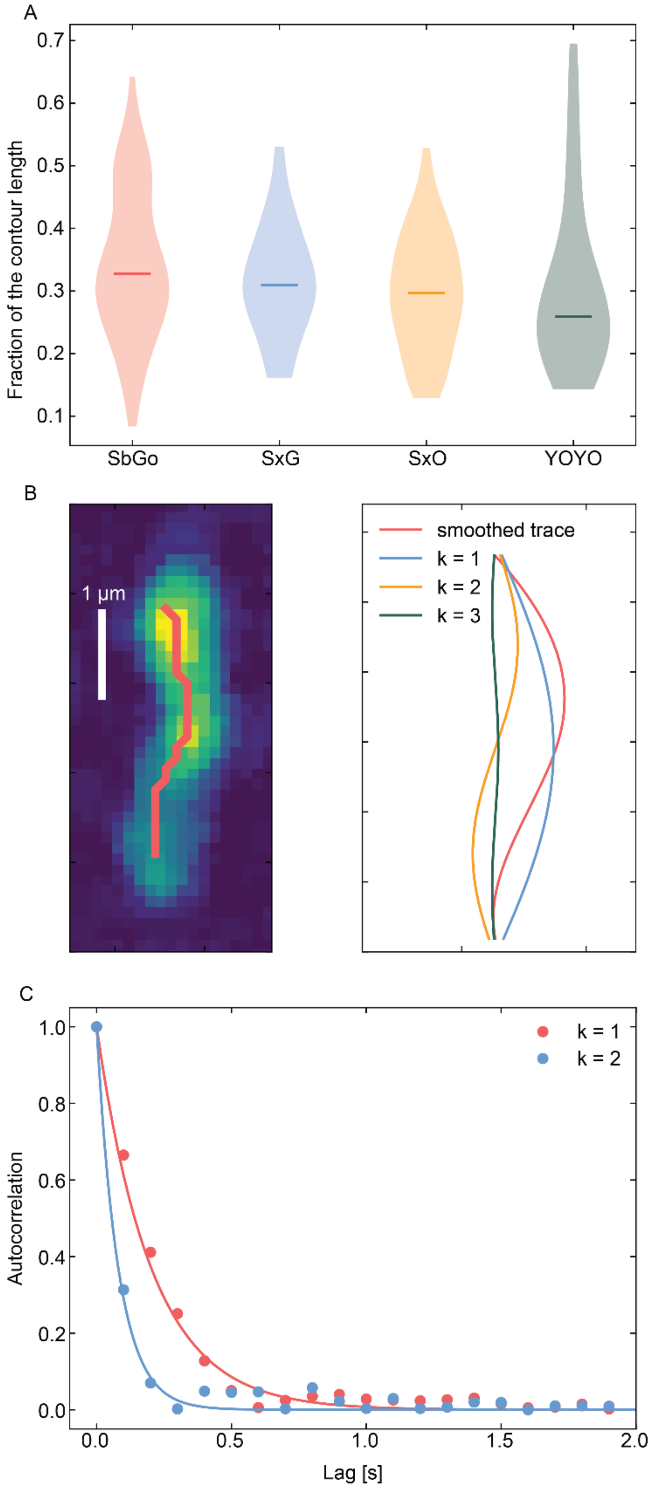


**Fig. S2. Measuring the persistence length from bending mode decomposition of intercalated DNA using single-molecule fluorescence imaging. A)** Distribution of DNA extension in the flow channel, normalized against the intercalated contour length. **B)** Representative example of 2.5 nM SYBR-Gold-labelled 50 kb DNA with its skeleton outline (left, skeleton in red) and its first three bending modes (right). **C)** Representative example of the autocorrelation function of bending mode coefficients $a_k$ for k= {1, 2}. Autocorrelation datapoints are represented as dots and exponential fits are plotted as lines in the corresponding colour. Exponential fits yielded time constants of 204 ms and 81 ms for k=1 and k=2, respectively.

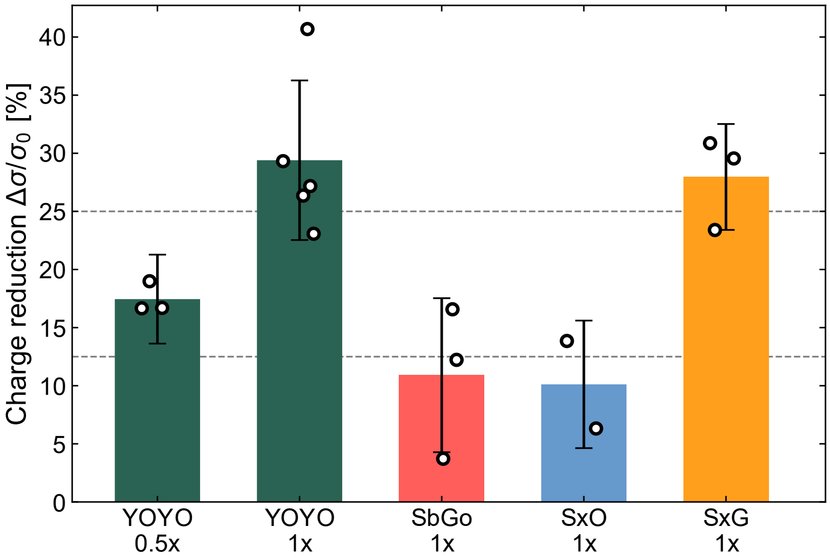


**Fig. S3. Surface charge reduction by intercalating dyes.** The reduction in DNA surface charge density calculated from the zeta-potential measurements as presented in Main Fig. 1F. The fractional reduction of the surface charge is calculated from the Grahame equation: $\Delta\sigma/\sigma_0 = 1 - \left[sinh\left(\frac{e\zeta}{2kT}\right)/sinh\left(\frac{e\zeta_0}{2kT}\right)\right]$. Here, $\zeta_0$ is the zeta potential of bare DNA (-42.6 ± 1.6 mV), and $\zeta$ is that of DNA stained with intercalating dye as reported in the main text and Fig. 1F. Error bars are standard deviations propagated from the uncertainties in both zeta potentials. Circles represent individual measurements. Dashed lines mark 12.5% and 25% reference charge neutralizations.

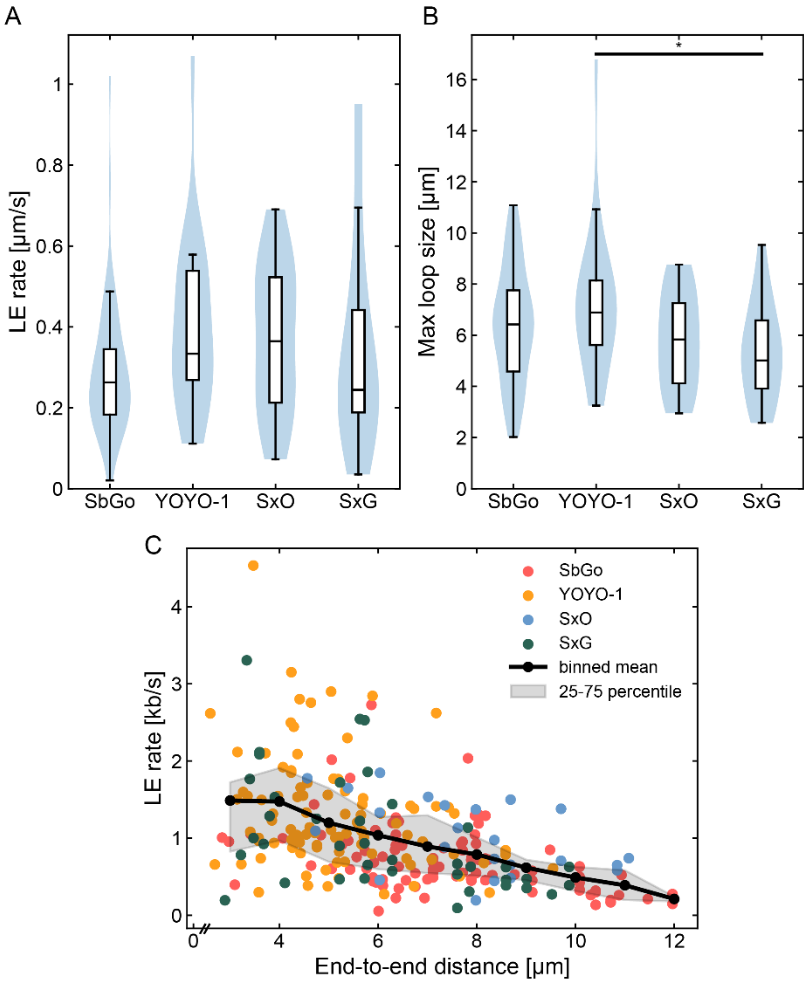


**Fig. S4. Loop extrusion on intercalated DNA.** These figures are related to main fig. 2B-D. Intercalators were present at a 10% binding density. **A)** Loop extrusion rate expressed in μm/s rather than kb/s. Note that we typically report the loop extrusion rate and loop sizes in base pair units rather than the contour length of base pairs and bound intercalators together. However, because intercalator binding lengthens the total contour of intercalated DNA – which scales with the intercalator density – the total amount of looped contour is a relevant value to report as well. **B)** Maximum loop size expressed in μm contour rather than kb. * indicates significant difference (Kolmogorov-Smirnov $p<0.05$). **C)** Loop extrusion rate in kb/s against the end-to-end distance of the whole molecule. Individual measurements are represented by filled circles. The black line represents the mean over all dyes, binned around each integer end-to-end distance (i.e. 3, 4, 5 μm, etc. ). The $25^{th}$-$75^{th}$ percentile is indicated by the shaded area. The downward slope of this line indicates that the loop extrusion rate goes down as the DNA molecule is stretched more. The mean loop extrusion rate decreases from 1.5 to 0.2 kb/s over the total range (3-12 μm).

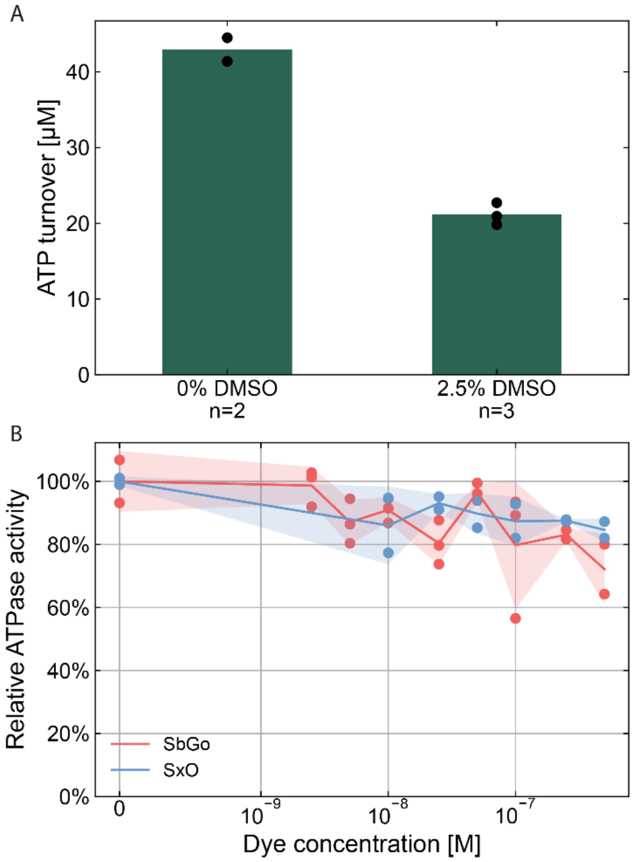


**Fig. S5. ATPase assay and the effect of DMSO and absolute concentration of intercalator. A)** The effect of DMSO at 2.5 % v/v on condensin. The ATPase is reduced by roughly 50% on the addition of an excess of DMSO. **B)** To quantify the effect of a high concentration of intercalator, we added similar absolute concentrations of intercalator as in Fig. 2E but in the presence of an excess of DNA such that the total intercalator density is <1% in all cases. As there is only a minimal effect on the ATPase activity, we conclude that the presence of intercalators itself has little contribution on the inhibitory effect, but rather it is the high binding density on DNA that inhibits condensin.

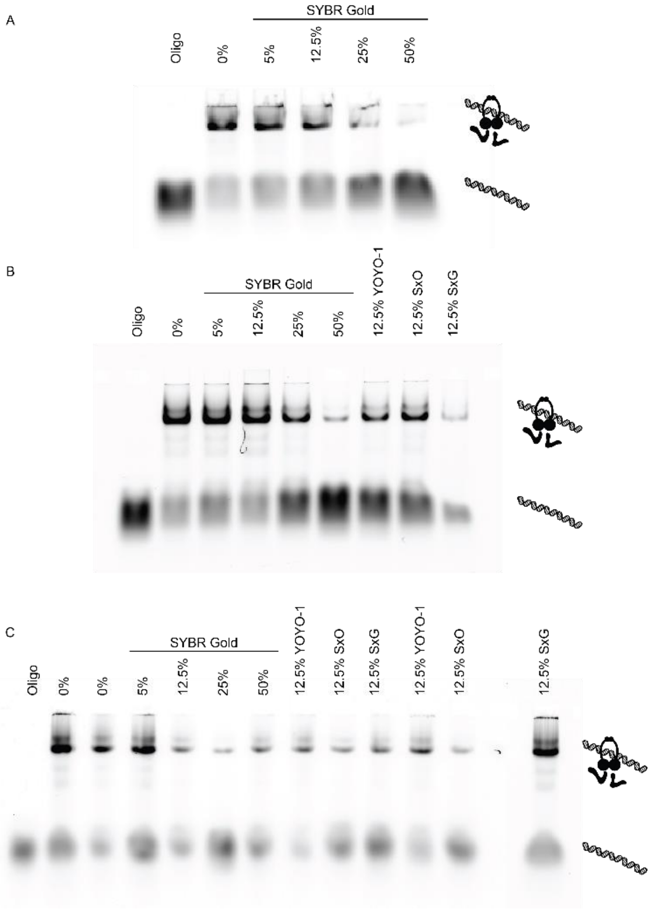


**Fig. S6. EMSAs of condensin binding to intercalated DNA. A-C)** Gel images of Cy5-labeled 42bp dsDNA in the presence and/or absence of condensin and/or intercalator. The conditions per lane are indicated above the lane, where all lanes except for the ones marked by "Oligo" contain oligo and condensin. The percentages indicate the intercalator binding density, such that 0% is only condensin and oligo DNA. The cartoons on the right-hand side represent the interpretation of the different bands: free oligo or condensin-bound oligo. We did not make a distinction between different binding modalities or stoichiometries. Lane quantifications are represented in main Figs. 2F-G.

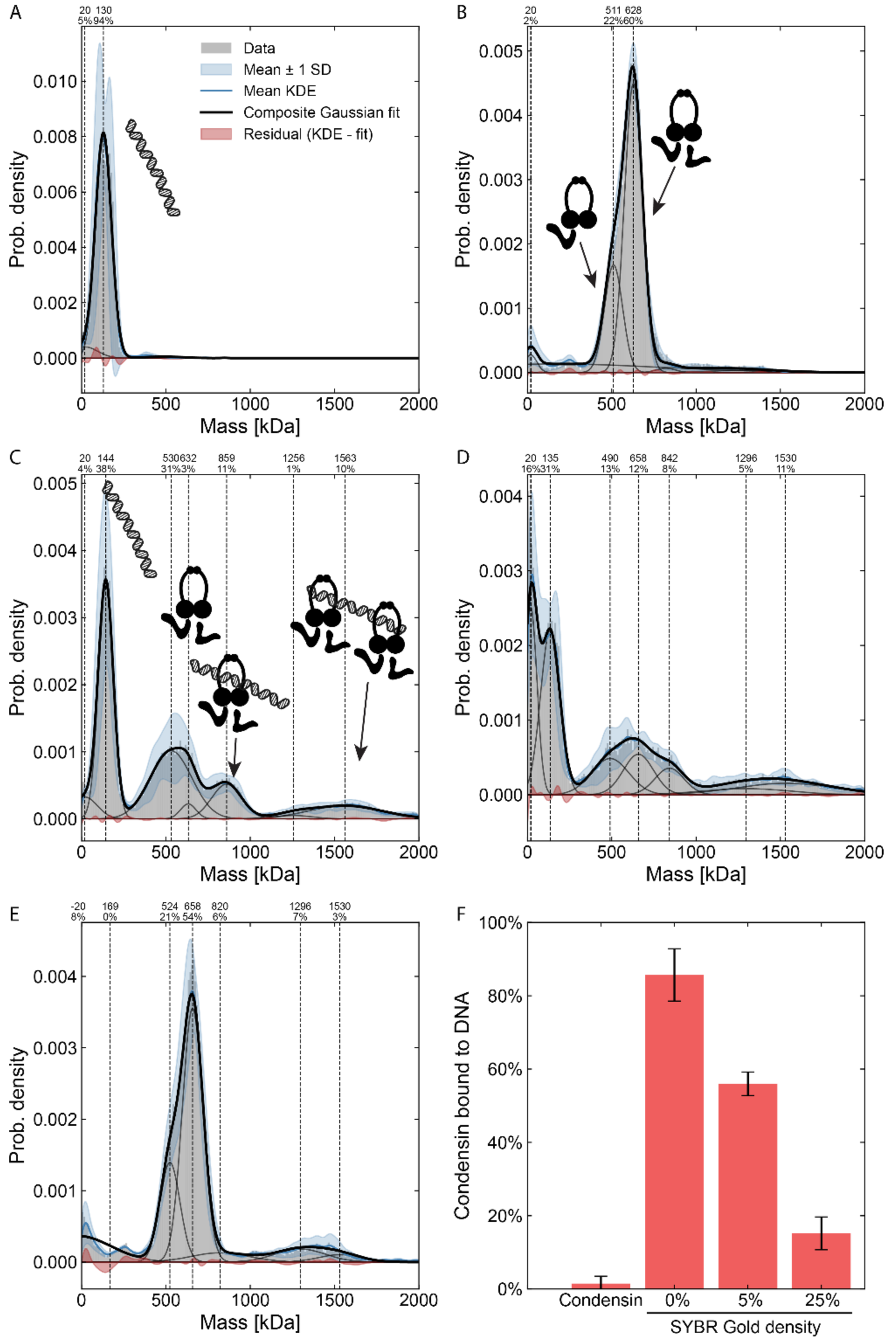


**Fig. S7. Mass photometry of condensin binding to DNA in LE buffer. A-E)** Mass photometry data. Masses were calibrated from mass photometry contrast using a known protein calibration, meaning that DNA will be observed at different masses than expected because of the different contrast-to-mass ratio for DNA and protein. Masses are depicted in the histogram shown in grey, while standard deviation of different replicates is represented as the blue shaded area. The mean kernel density estimate (KDE) with a bandwidth of 20 kDa is generated from the replicates, which is then used to fit Gaussian peaks at each known peak position. Gaussian peaks are represented as dark grey lines and the difference between the mean KDE and Gaussian fitted peaks is shown in red. Peak positions are called within 20 kDa of the reference peak position. **A)** 500 bp DNA is observed around 130 kDa; N = 3. **B)** Condensin is observed in two peaks. The largest peak is positioned at 628 kDa, which is close to the expected mass of the complex based on sequence, whereas a smaller peak is observed at 511 kDa. We attribute the smaller peak to a population of condensin complexes that lack the Ycg1 subunit, for which the expression is under cell-cycle control. This small population is not observed to participate in DNA binding in the subsequent panels. N=4. **C)** Condensin binding to DNA. 86% of all intact condensin complexes were observed to bind to DNA in either of two peaks: 859 kDa and 1563 kDa. The first peak is attributed to 1:1 condensin:DNA, whereas the second peak is attributed to 2:1 condensin:DNA. The contribution of DNA to the total mass is also seen to increase upon binding, indicating that the contrast-to-mass ratio changes upon protein binding. N=6. **D)** Condensin similarly binds to DNA in the presence of 250 nM SYBR Gold (5% intercalator density on DNA). N=4. **E)** Condensin binding to DNA in the presence of 1.26 μM SYBR Gold (25% intercalator density on DNA). N=3. **F)** Quantification of the fractions of condensin bound to DNA in panels B-E. In the presence of increasing amounts of intercalator, the DNA-bound peaks decrease and the condensin-alone peak increases, consistent with condensin lowering its affinity to intercalated DNA. Panel B serves as a reference for the background of condensin complex found around the expected peak positions of condensin bound to DNA, which yields only 1.5% background. Error bars represent the standard deviation of the bound fractions, calculated based on the delta method for the ratio of means over each replicate and the ratio of peak fractions. The single-shot $K_d$ for condensin binding to DNA was calculated for these conditions: 5.3 nM [2.1, 9.2], 27.1 nM [22.9, 31.9], 214.6 nM [142.1, 378.2] (1 standard deviation confidence interval in square brackets) for 0%, 5%, and 25% SYBR gold intercalator density, respectively.

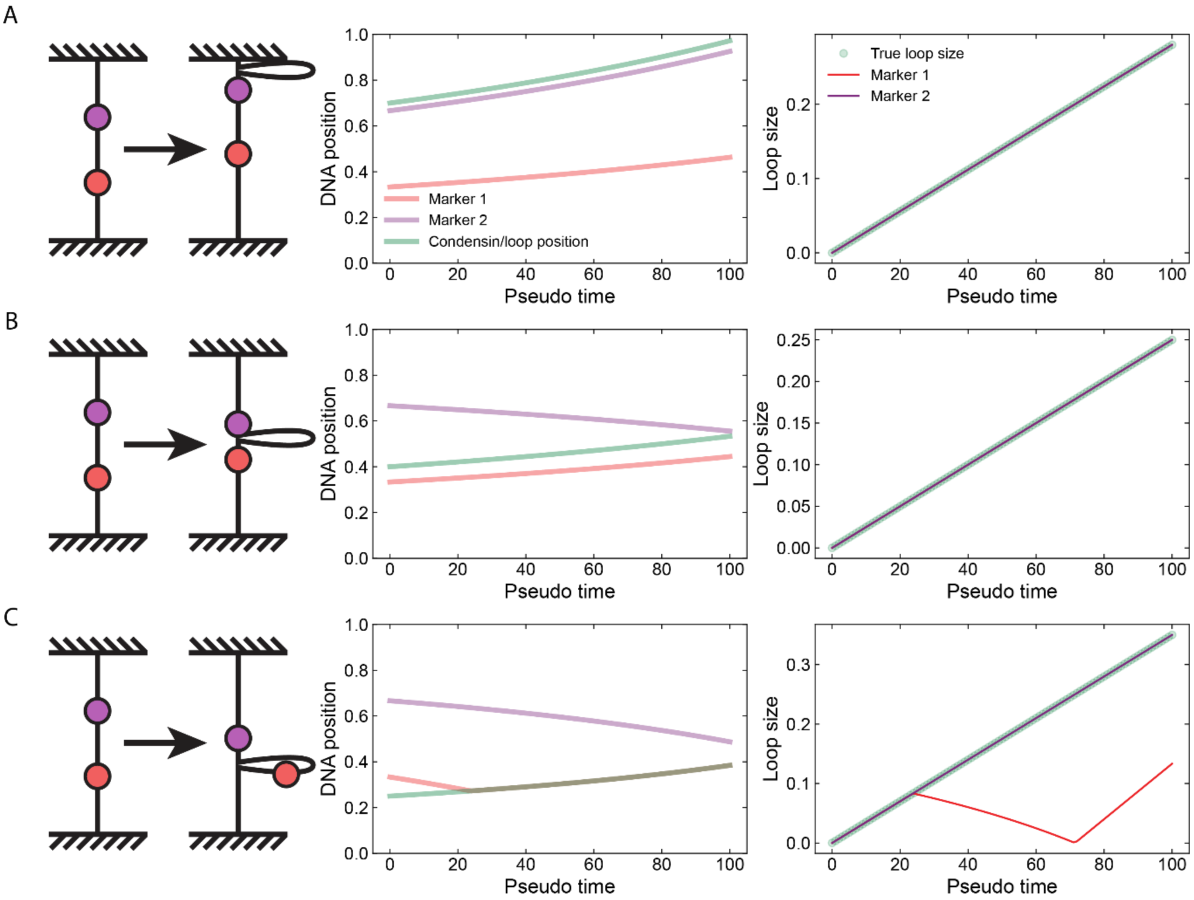


**Fig. S8. Modified loop extrusion assay based on spot label markers in three different scenarios.** Simulations were run to model the positions of spot label markers in relation to the loop size and condensin. Sketch of the scenario is depicted on the left, positions on DNA are plotted in the middle, and the inferred loop size and the true loop size are plotted on the right. **A)** Scenario as depicted in Fig. 3A-E: A loop starts at marker 2 and moves upward toward the end of the DNA, taking along both markers. The loop size can be calculated accurately from both marker positions. **B)** Scenario as depicted in Fig. 4A: A loop starts in the middle between the two markers and draws both markers in. The loop size can be calculated accurately from both marker positions in this scenario, too. **C)** A third common scenario: the loop is extruded towards a marker and passes over the marker, taking it along with it. Because the loop was moving in the direction of this marker and continues in the same direction, the marker is moved back towards its original position (visible from timepoint 20 to 70), which looks like a decrease in the loop size until the loop is returned to its origin. If the loop continues onward in the same direction, the marker would indicate an increase in loop size again (visible from timepoint 70 on). Because of this tell-tale pattern, the correct loop size can be inferred from the unlooped marker.

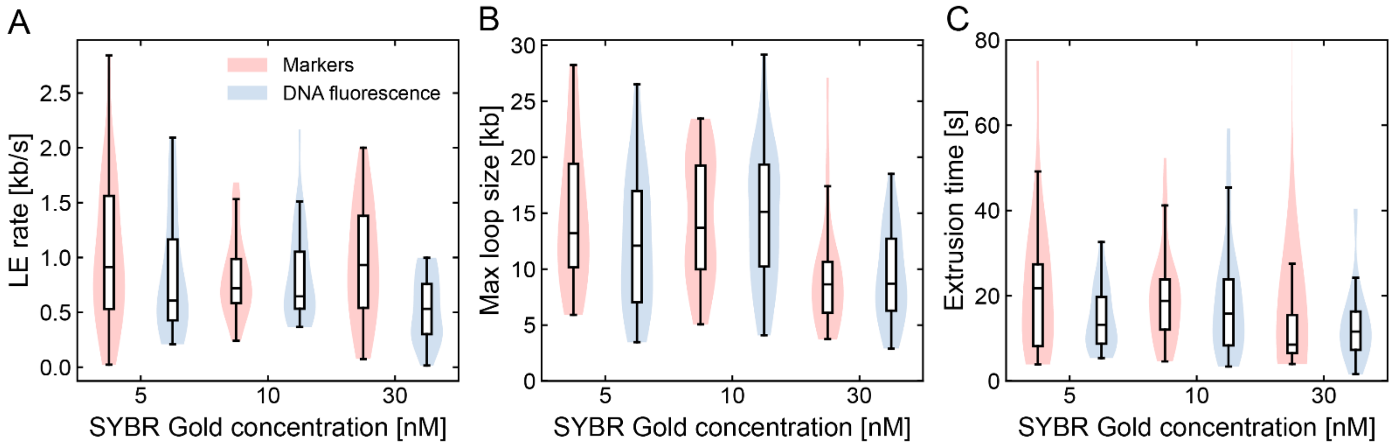


**Fig. S9. Loop extrusion properties from markers and DNA fluorescence compared.** The datapoints are the same as in main Fig. 4B-D, but now the loop extrusion properties are calculated from the marker spot labels and DNA fluorescence intensity and compared side-by-side. **A)** Loop extrusion rate, **B)** maximum loop size, and **C)** Loop extrusion time. As is clear, the properties calculated from fluorescence intensity qualitatively similar, but quantitatively different, except for the maximum loop size, which is the same between the two methods.

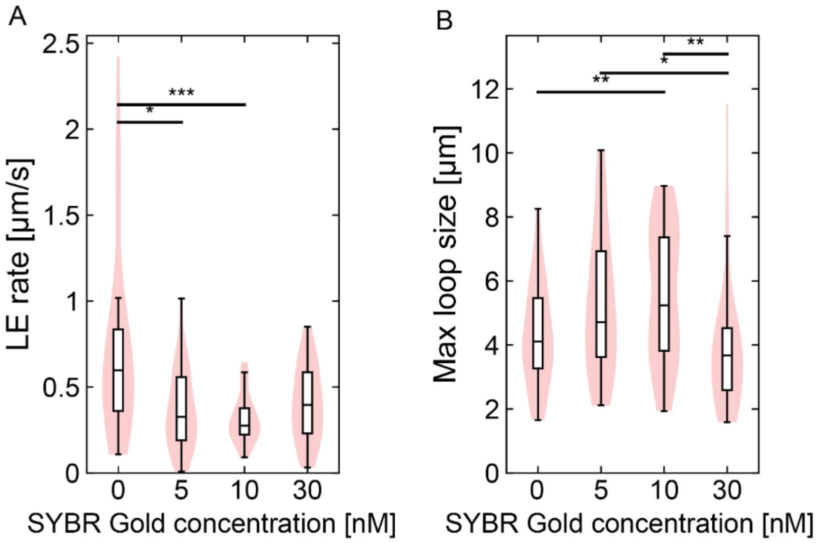


**Fig. S10. Loop extrusion quantification based on DNA contour rather than basepairs related to main Fig. 4B-D. A)** Loop extrusion rate expressed in contour length of the DNA-intercalator complex per second rather than kb/s. **B)** Maximum loop size expressed in contour length, similar to A. As intercalators add contour length to the DNA upon binding, rate and loop size are higher for higher intercalator concentrations when expressed as total contour length. Significant differences (Kolmogorov-Smirnov) are indicated as: *: $p < 0.05$, **: $p < 0.01$, ***: $p < 0.001$. N= 39, 25, 36, 27, respectively.
